# Branched-chain keto acids promote an immune-suppressive and neurodegenerative microenvironment in leptomeningeal disease

**DOI:** 10.1101/2023.12.18.572239

**Authors:** Mariam Lotfy Khaled, Yuan Ren, Ronak Kundalia, Hasan Alhaddad, Zhihua Chen, Gerald C. Wallace, Brittany Evernden, Oscar E. Ospina, MacLean Hall, Min Liu, Lancia N.F. Darville, Victoria Izumi, Y. Ann Chen, Shari Pilon-Thomas, Paul A. Stewart, John M. Koomen, Salvatore A. Corallo, Michael D. Jain, Timothy J. Robinson, Fredrick L. Locke, Peter A. Forsyth, Inna Smalley

## Abstract

Leptomeningeal disease (LMD) occurs when tumors seed into the leptomeningeal space and cerebrospinal fluid (CSF), leading to severe neurological deterioration and poor survival outcomes. We utilized comprehensive multi-omics analyses of CSF from patients with lymphoma LMD to demonstrate an immunosuppressive cellular microenvironment and identified dysregulations in proteins and lipids indicating neurodegenerative processes. Strikingly, we found a significant accumulation of toxic branched-chain keto acids (BCKA) in the CSF of patients with LMD. The BCKA accumulation was found to be a pan-cancer occurrence, evident in lymphoma, breast cancer, and melanoma LMD patients. Functionally, BCKA disrupted the viability and function of endogenous T lymphocytes, chimeric antigen receptor (CAR) T cells, neurons, and meningeal cells. Treatment of LMD mice with BCKA-reducing sodium phenylbutyrate significantly improved neurological function, survival outcomes, and efficacy of anti-CD19 CAR T cell therapy. This is the first report of BCKA accumulation in LMD and provides preclinical evidence that targeting these toxic metabolites improves outcomes.

---

Leptomeningeal disease (LMD) is a rapidly progressing and dreaded complication of cancer that is clinically detected in 5-15% of patients with late-stage solid tumors ^1^ and is apparent at autopsy in up to 20-30% of cancer patients with metastatic disease and neurologic symptoms ^2–4^. LMD occurs when malignant cells seed to the leptomeninges (membrane coverings of the brain and spinal cord) and cerebral spinal fluid (CSF) compartments ^5^. Rapid and debilitating neurological symptoms drastically impact LMD patients’ survival outcomes. The median survival time for treated LMD patients is 2–6 months, with death typically occurring due to progressive neurologic dysfunction ^6–11^. For the majority of the patients, the condition is rapidly terminal, and the primary goal of treatment is to improve patients’ neurological function and quality of life through palliative and supportive care. Current LMD treatments include combination radiotherapy regimens with intrathecal chemotherapy (ex: cytarabine, methotrexate, thiotepa, etc) or intrathecal targeted/immune therapy ^12–14^. However, studies have shown variable/limited responses to treatment, which are frequently further complicated by the toxicity of the regimens ^14,15^. Systemic therapy can be added to control extracranial disease and potentially prolong patient survival ^14^.

Non-Hodgkin B cell lymphomas (NHBCL) comprise 90% of all malignant lymphoma cases. For these patients, LMD may manifest either as metastatic dissemination of extracranial disease (occurring in 5-10% of NHBCL patients) ^16^ or as the primary disease site, as in the case of 7-40% of primary central nervous system lymphomas^17,18^. NHBCL, typically a systemic disease, is often well-controlled with combination chemo-immunotherapy. CNS recurrence, although uncommon, happens rapidly and frequently as the sole site of relapse when it occurs ^19^. Multiple studies have now demonstrated that CD19-targeting CAR T-cell therapies are effective at treating chemo-refractory NHBCL; however, the anti-tumor effects on responding LMD tumors are rarely long-lasting^20–24^, despite clear evidence of CAR T-cell penetration into the CSF space^25^. Therefore, there is an urgent clinical need to better understand the biology of this understudied anatomical site that promotes tumor growth, drug resistance, and neurological deterioration manifested in LMD patients. In this study, we implemented a multifaceted strategy to characterize the proteomic, metabolic, and immune environment within LMD using patient specimens, *in vivo* models, and *in vitro* functional studies. Our study provides the first preclinical evidence for combining branched-chain keto acid-lowering treatments with anti-CD19 CAR T cell therapies to improve the quality of life and survival outcomes for LMD patients.

## RESULTS

### The cellular landscape of leptomeningeal lymphoma

To define the cellular landscape of leptomeningeal lymphoma, we performed single-cell RNA sequencing (scRNAseq) analysis on eight CSF specimens from seven patients, including four CSF specimens from Non-Hodgkin’s B-cell lymphoma patients with leptomeningeal involvement (**Supplemental Table 1**). Of the LMD specimens, two serial samples came from a rare patient who responded well to intrathecal chemotherapy and had a long overall survival time of over 56 months post-LMD diagnosis. Two additional samples were from two patients with a more typical survival time of <1-4 months post-LMD diagnosis despite aggressive therapy (**Supplemental Figure 1**). Four CSF specimens from patients with no history of malignancy were included as controls (**Figure 1A**). Cell profiling analysis identified seven broad cell categories within the CSF, including B cells, CD4 T cells, CD8 T cells, natural killer (NK) cells, macrophages, monocytes, and plasmacytoid dendritic cells (**Figure 1B**). There was a substantial overlap of cell clusters from various samples, especially non-tumor clusters, suggesting that cell clustering is primarily driven by cell type instead of the sample of origin (**Supplemental Figure 2A**). The total number of cells in CSF of patients without LMD is minimal (<300 in 7mL) and largely consists of CD4 T cells (**Figure 1C-D, Supplemental Figure 2B**). Meanwhile, LMD CSF shows the infiltration of tumor and immune cells (**Figure 1C-D, Supplemental Figure 2A**). The CSF of these patients shows a considerable accumulation of B cells, as expected from B cell lymphoma tumors, making up approximately half of the cellular material in CSF (**Figure 1C-D, Supplemental Figure 2B**). Looking closer at the non-B cell lymphoid landscape, we identified 11 clusters of T and NK cells, including 3 clusters of CD4 T cells, 7 clusters of CD8 T cells, and one cluster of NK. (**Figure 1E, Supplemental Table 2**). The number of T cells identified in CSF of patients positive for leptomeningeal disease was much greater than those negative for the disease (**Supplemental Figure 2C**).

**Figure 1:**
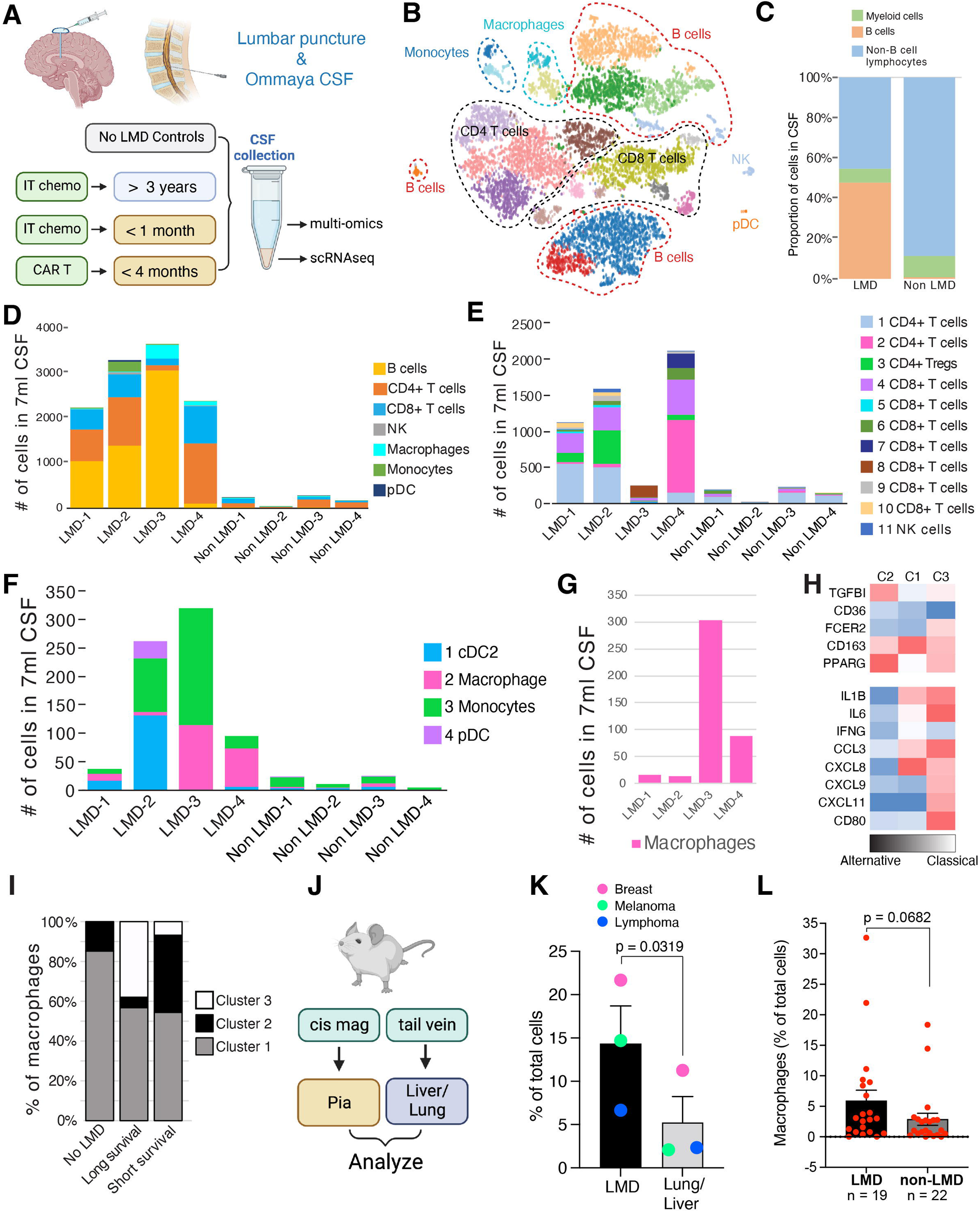
The cellular landscape of leptomeningeal lymphoma. A. Schematic illustration demonstrating the CSF sampling from lymphoma-LMD patients, treatments received at the time of collection, and the patient’s survival time. CSF from non-LMD were also collected as controls. B. Uniform manifold approximation and projection (UMAP) plot of all cell populations in all CSF specimens. C. The proportion of myeloid cells, B cells, and non-B cell lymphocytes in lymphoma LMD versus non-LMD CSF samples. D. The number of cells from the main cell types in 7mL of lymphoma-LMD and non-LMD CSF. E. The number of T and NK cells from each of the 11 clusters identified in each 7mL of lymphoma-LMD and non-LMD CSF. F. The number of myeloid cells from each of the 4 clusters identified in 7mL of lymphoma LMD and non-LMD CSF. G. The number of macrophages in each 7mL CSF sample from lymphoma LMD. H. Expression of key markers of different macrophage activation states to classify macrophage subclusters into C1, C2, and C3. I. Percentage of each macrophage subcluster (identified in H) according to prognosis. J. Outline of the cell injections and sample collection for single-cell RNAseq of tissues from animal models of LMD. K. Macrophages as a percentage of total cells identified using scRNAseq of leptomeningeal layer vs. extracranial metastatic mouse tissues from each of the breast, melanoma, and lymphoma models. Bars represent mean ± SEM. L. Bar graph showing the abundance of macrophages (as a percentage of total cells) identified by scRNAseq in CSF samples from melanoma LMD patients versus tissues from non-LMD tumor sites of melanoma patients.

In the myeloid cell lineages, we identified four major clusters of cells, including classical dendritic cells 2 (cDC2), plasmacytoid dendritic cells (pDC), macrophages, and monocytes (**Figure 1F, Supplemental Table 3**). Similar to the lymphoid compartment, the number of myeloid cells identified in CSF of patients positive for leptomeningeal disease was much greater than those negative for the disease (**Supplemental Figure 2D**). Samples from patients with short survival (LMD-3 and LMD-4) show an accumulation of macrophages and a lack of antigen-presenting dendritic cells in the CSF space: features not observed in the extraordinary survivor (**Figure 1G**). A closer examination of macrophages identified three major clusters based on overall gene expression profiles (**Figure 1H**). Among these three groups, we observed differences in the expression of markers associated with the pro-inflammatory classical polarization and pro-tumorigenic alternative polarization (**Figure 1H**). We noted that cluster 2 had the highest expression of alternative activation markers such as TGFB1, PPARG, and CD163 and expressed no classical activation markers, whereas cluster 3 exhibited an increased expression of classical activation markers. Among the patients, we observed the highest enrichment of alternatively activated cluster 2 in the patients with short survival and the greatest number of classically activated macrophages in the patient with prolonged survival (**Figure 1I**). Next, we validated the LMD-specific accumulation of macrophages in murine models of LMD. A20 lymphoma, 4T1 breast cancer, and SM1 melanoma cell lines were injected intrathecally into the CSF space or via the tail vein (**Figure 1J**). ScRNAseq analysis of pia, livers, and lungs confirms that the accumulation of macrophages is more profound in the leptomeningeal disease and is not limited to LMD from lymphoma but instead suggests a pan-cancer phenomenon (**Figure 1K**). Analysis of our previously published melanoma scRNAseq cohort consisting of 8 skin metastasis biopsies, 14 surgical brain metastases, and 19 CSF specimens from patients with LMD also confirms a trend (p=0.0682) for increased macrophage accumulation in LMD compared to brain parenchyma metastases or extracranial skin metastases (**Figure 1L**)^26^.

### The immune-suppressive cellular microenvironment of lymphoma LMD

To understand the landscape of lymphocytes in LMD in finer detail, we examined gene expression profiles associated with activation, proliferation, exhaustion, anergy, senescence, regulatory T cells, and naïve T cells across the 11 clusters of T/NK cells and assigned them to putative functional groups (**Figure 2A**). Interestingly, the T cell landscapes between the samples from the patient with prolonged survival and from those with short survival show distinct T cell landscapes. Both CSFs from the patient with long survival show large proportions of cluster 1 CD4 T helper cells and cluster 3 CD4 regulatory T cells (**Figure 2B**). Significant T-cell infiltration was noted in sample LMD-4 from a patient treated with CART, particularly from clusters 2, 4, 6, and 7 (**Figure 1E**). While the CSF from the extraordinary survivor showed multiple (albeit small) T cell clusters that are active and proliferating as well as those that are active approaching exhaustion, the majority of T-cells in the CSF of patients with short survival were naïve, anergic, exhausted or senescent, with no active, proliferating T cell clusters present (**Figure 2B**). Not surprisingly, we found very few T cells in the CSF of LMD-negative patients (**Supplemental Figure 2C**). Of the T cells present in LMD-negative specimens, the majority were naïve (**Figure 2B**).

**Figure 2.**
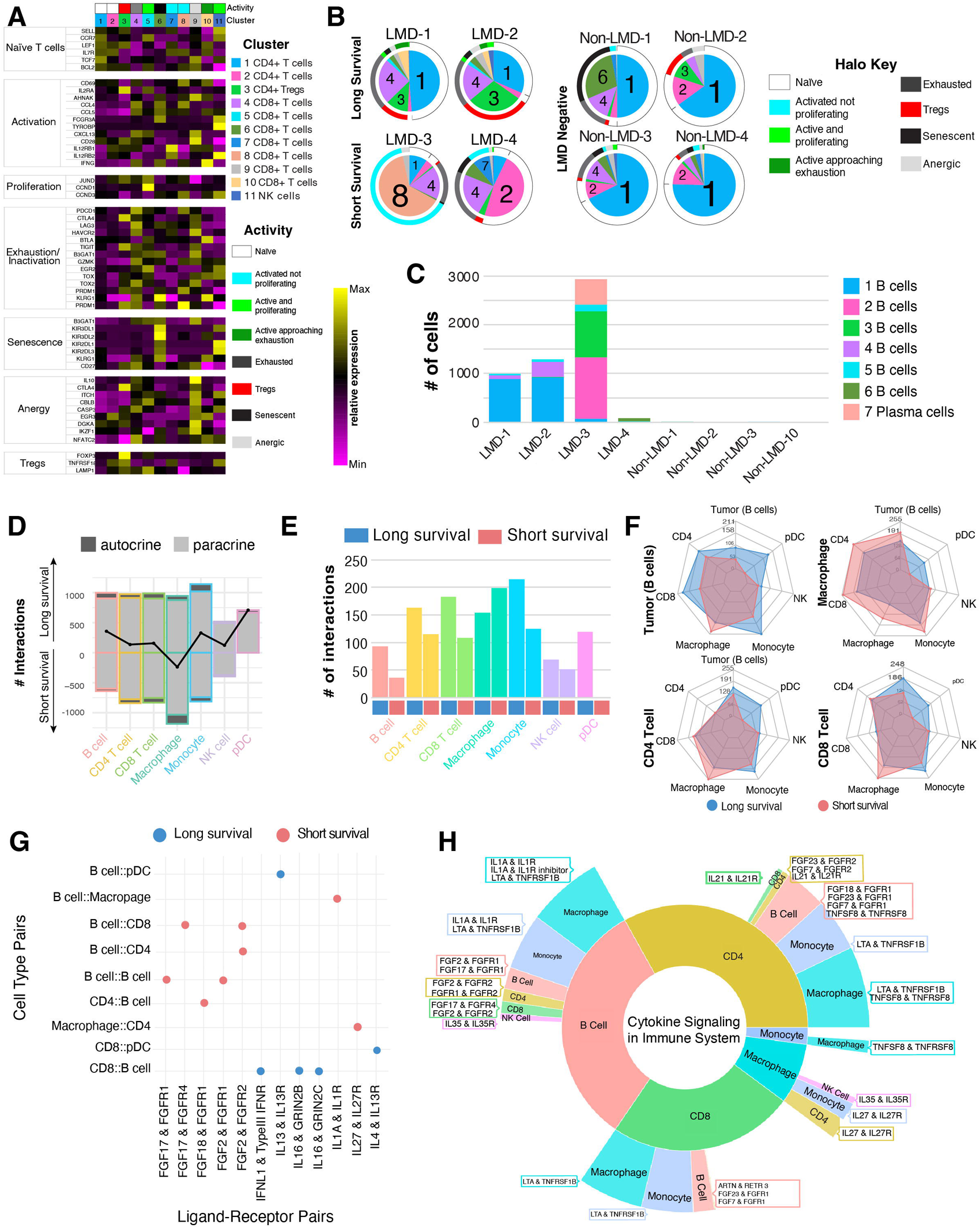
The cellular microenvironment of lymphoma LMD is immune-suppressive. A. Heat map showing the expression of key markers used to assign functional/activation groups to each of the 11 T and NK cell clusters. B. Pie charts show the distribution of each T/NK cluster in lymphoma LMD and non-LMD samples. The colored halo indicates the predicted functional status of each T/NK cell based on gene expression profiles. C. The number of cells found in each of the seven subpopulations of B cells in 7mL of CSF from lymphoma LMD and non-LMD patients. D. Back-to-back bar plots show the direction of interaction per cell type, comparing the patients with a long survival (positive axis) vs. short survival (negative axis). E. The bar plot shows the number of interactions per cell type in the patients with long survival and short survival. F. Radar plot demonstrating the communication between a specific cell type and all other cell types, comparing the patients with long survival (blue) vs. short survival (red). G. Dot blot for selected unique ligand-receptor pairs to each condition. All the unique ligand-receptor pairs for different cell type pairs are shown in Supplementary Tables 6 and 8. H. Sunburst plot for cytokine signaling pathways as the most significant functional term based on only the unique ligand-receptor in patients with short survival. The width of each section represents the relative fraction of interactions (weighted by score) enriched in that cell type. Boxes show specific int-pairs enriched for the corresponding cluster pairs. All unique, condition-specific functional terms are shown in Supplemental Tables 5 & 7. Plots in D, E, F, G, and H were generated in InterCellar’s multiple conditions module.

To better understand the tumor cell landscape of LMD, we took a closer look at B cell heterogeneity. We identified six B cell clusters and one plasma cell cluster (**Figure 2C**). Interestingly, the principal patient-driven heterogeneity was observed in the tumor cells, where each patient demonstrated a specific landscape of predominant cell clusters, with minor overlap in clusters 1 and 5 among multiple patients (**Figure 2C**). Gene expression profiles distinguishing each cluster were further analyzed (**Supplemental Table 4**). The dominant cluster of B cells from LMD-1 and LMD-2 samples was cluster 1 and expressed high levels of CCND1 and Sox11, consistent with the distinctive profile of cyclin D positive mantle cell lymphoma ^27,28^. Interestingly, this long-surviving patient also had a subpopulation of B cells (cluster 4) expressing high FOSL2, a known mediator of autoimmunity and inflammation ^29^ (**Supplemental Figure 3**). The two dominant B cell populations in sample LMD-3 from a patient with short survival, clusters 2 and 3, expressed high levels of ARGLU1, a known repressor of interferon type 1 signaling ^30^ (**Supplemental Figure 3**). Finally, LMD-4 mostly contained cluster 7 B cells expressing high levels of TCL1B and INHBA, markers associated with high-grade leukemias and lymphomas^31,32^. Importantly, no significant difference was observed in the expression of CD19 or CD20 in the B cell clusters, which are the common targets of anti-lymphoma CART cells ^33,34^ (**Supplemental Figure 4**).

While comparing the numbers and proportions of cells and cell types among the samples provided us with an initial picture of the overall cellular composition in Lymphoma LMD, we wanted to learn more about how different subpopulations of cells may interact within these cellular landscapes. Therefore, we interrogated the cell-cell interactions within the cellular landscape of lymphoma LMD. Interestingly, we found fewer autocrine and paracrine B-cell, T-cell, and dendritic cell interactions in CSF of patients with short survival, whereas we saw more macrophage interactions in these patients (**Figure 2D-F**). There were also more interactions between macrophages and tumor cells, CD4 T cells, CD8 T cells, and monocytes (**Figure 2F**). Looking closer at the unique ligand-receptor interactions per each condition using CellPhoneDB with InterCellar visualization (see Methods) (**Figure 2G-H, Supplemental Figure 5, Supplemental Tables 5-8**), we found a prevalence of interactions involved in cytokine signaling in the patients with short survival, including signaling through fibroblast growth factor receptor (FGFR) and other inflammatory cytokines creating an immune suppressive TME (**Figure 2G**) ^35^. On the other hand, CD8 cells in the CSF of the patient with a long survival express IFNL1, known as IFNλ a type III interferon cytokine (**Figure 2G**). IFNλ is a potent antiviral cytokine in patients ^36^ and showed a beneficial anti-tumor activity in preclinical models ^37^. Interestingly, IL13R was uniquely expressed on the dendritic cells in the long survival, which may contribute to anti-tumor immune response within TME (**Figure 2G**). Dendritic cells pulsed only with IL13Rα2 peptides were investigated as an autologous vaccine in glioblastoma, but only a subgroup of patients showed a robust immune response (NCT01280552) ^38^. The most significant functional term based on the unique ligand-receptor interactions among cells in patients with short survival was cytokine signaling in the immune system (**Figure 2H**). Ligand-receptor pairs specific to this category highlight the interactions among cells mediated via cytokines and growth factors such as FGFR, IL-1α, and IL-27, which have immune-suppressive tumor-promoting functions in the TME ^39,40^. On the other hand, the most significant functional term for ligand-receptor interactions unique to patients with long survival was integrin signaling, which included numerous interactions among integrins and collagens (**Supplemental Figure 5**). Integrin engagement plays a major role in effective immune activation by promoting immune cell aggregation and TCR signaling^41,42^.

### Multi-omic analysis of patient CSF suggests an upregulation of processes involved in immune regulation and progressive neuronal degeneration

To examine how the unique LMD microenvironment may promote the immune-suppressive cellular landscape, we performed a comprehensive multi-omic analysis of the acellular CSF fluid from patients with and without lymphoma LMD. Overall, 27-32% of protein, metabolite, and lipid content was significantly altered in the CSF of patients with LMD (**Figure 3A, Supplemental Tables 9-11**). We noted that lipid classes involved in membrane composition, including phosphatidylethanolamine (PE), were significantly depleted in patients with LMD, consistent with an environment of a rapidly dividing tumor (**Figure 3B, C**). Furthermore, we observed a significant reduction of ceramide and prenol lipids, which are important in immune regulation ^43^. On the other hand, we noted a significant elevation in four lipids of the lysophosphatidylcholine class, which play a role in neuron demyelination (**Figure 3B, C**) ^44,45^. Along the same lines, of the few proteins significantly depleted from CSF in LMD patients, several are involved in critical processes of normal neuron function, including GRIA4 (learning and memory) ^46^, ARSA (essential enzyme regulating myelination) ^47,48^, THY1 (axon regeneration)^49^, CDH4 (neuronal outgrowth)^50^, NPTXR (synaptic activity) ^51^, neurotransmitters TAC1 and TAC3 (behavioral responses) (**Figure 3D**) ^52^. Principal component analysis (PCA) of proteomic and lipidomic profiling highlights the separation of samples from patients with short survival (LMD-3 and LMD-4) from the rare patient with long survival (LMD-1 and LMD-2) and from the non-LMD controls (**Supplemental Figure 6A-B**). Pathway enrichment analysis of differentially expressed proteins highlighted processes associated with innate immune responses (including macrophage function and immunodeficiency) and neurodegenerative disorders, including Alzheimer’s and multiple sclerosis (**Figure 3E, Supplemental Table 12**). With these strong suggestions of neurodegenerative processes, we specifically examined the abundance of proteins whose accumulation or depletion was associated with neurodegenerative diseases such as Alzheimer’s and Parkinson’s ^53–55^. Immunoglobulin G, transferrin, vitamin-D binding protein, PARK7, and DJ-1 were significantly enriched in CSF from LMD patients (**Figure 3F**). These same CSF samples were also significantly depleted of amyloid β precursor, neurosin, and BDNF (**Figure 3F**). The signatures associated with neurodegenerative processes were also present in our previous large cohort of serial CSF specimens from patients with melanoma LMD (**Supplemental Table 13**) ^56^. The enrichment and depletion of many of these markers were statistically significant over the course of the patients’ LMD progression (Spearman correlation, **Supplemental Table 13**). A closer comparison of the LMD-specific proteome demonstrates a high level of agreement among proteins with the greatest enrichment (primarily those involved in innate immune response) and depletion (mostly those involved in normal neuronal function) in the lymphoma and the melanoma LMD cohorts compared to no LMD controls (**Figure 3G**). Taken together, these data suggested an immune-suppressed, neurodegenerative tumor microenvironment in LMD but did not identify the mechanistic driver of this environment.

**Figure 3.**
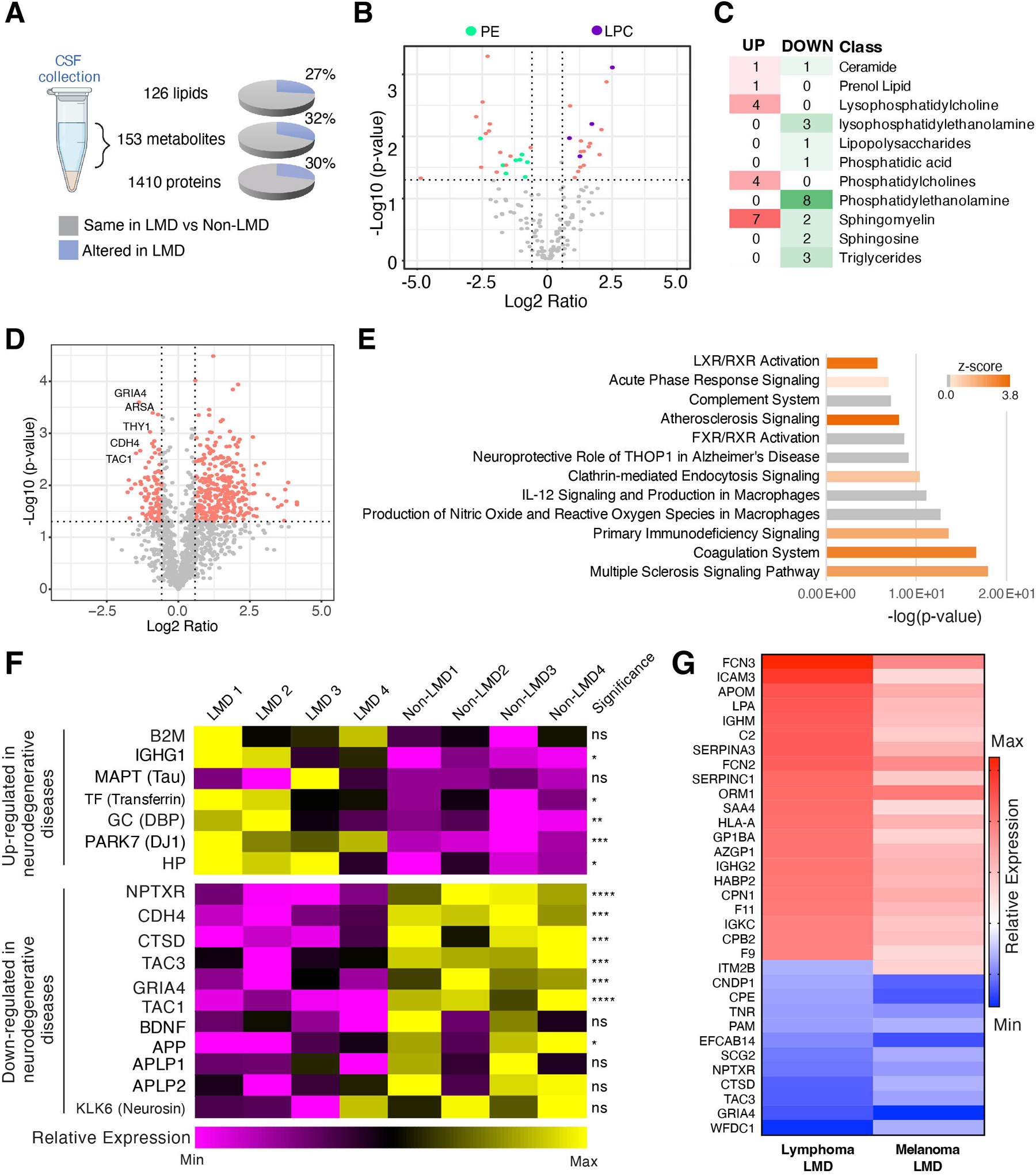
Proteomic and lipidomic characterization of LMD tumor microenvironment. A. Schematic illustration outlining the multi-omic analysis and the number of identified lipids, metabolites, and proteins in the CSF. Percentages indicate differential expression between lymphoma-LMD and non-LMD controls. B. Volcano plots show differentially abundant lipids in the CSF of lymphoma patients with LMD compared to no LMD controls. The significantly altered abundance of lipids in the lysophosphatidylcholine (LPC) lipid class is denoted in purple, and the significantly altered abundance of lipids in the phosphatidylethanolamine (PE) lipid class is denoted in green. C. The number of lipids identified to have significant changes in abundance in lymphoma LMD compared to non-LMD control for each lipid class. D. Volcano plots show differentially abundant proteins in the CSF of lymphoma patients with LMD compared to non-LMD controls. E. Pathway enrichment analysis of the differentially expressed proteins using Ingenuity Pathway Analysis. F. Heatmap showing the individual relative expression of neurodegenerative disease-associated proteins in lymphoma-LMD and non-LMD. Statistical significance was assessed using Welch’s t-test. G. Heat map showing the top upregulated and downregulated proteins in CSF from lymphoma LMD and their expression in melanoma LMD, highlighting the equivalent processes of complement activation and neurodegeneration in LMD regardless of the primary tumor type.

### Metabolomic analysis of patient CSF uncovers a dramatic accumulation of branched-chain keto acids

Since the metabolic environment of CSF is unique compared to other common sites of metastatic disease, we sought to better understand the metabolic microenvironment of LMD using untargeted metabolomic profiling of the patients’ CSF. Again, the PCA scores plot highlights the separation of LMD3 and LMD4 samples from patients with short survival from LMD2 and LMD1 from the extraordinary survivor and the non-LMD controls (**Supplemental Figure 6C**). Integration of the proteomic and metabolomic data shows the most significant alterations in complement and coagulation cascades, cell adhesion processes, and multiple metabolic processes, including central carbon metabolism, citrate cycle, pentose phosphate pathway, pyruvate metabolism, and glutathione metabolism, emphasizing a very broad range of metabolic alterations in the CSF microenvironment with LMD (**Supplemental Figure 6D**). Analysis of individual differentially abundant metabolites identified a striking accumulation of branched-chain keto acids (BCKA) α-ketoisocaproate (KIC)/α-keto-β-methylvalerate (KMV) in CSF of LMD patients (**Figure 4A**). Branched-chain keto acids are abnormal metabolites that result from an incomplete metabolism of branched-chain amino acids^57^ and are known as metabotoxins, neurotoxins, and acidogens. BCKAs are typically associated with a group of severe metabolic disorders termed the Maple Syrup Urine Disease, where BCKA and branched-chain amino acid accumulation are linked to neurological dysfunction^58,59^. Process enrichment analysis of differentially abundant (p <0.05) metabolites showed branched-chain amino acid biosynthesis and degradation pathways to be enriched in patients with LMD, and we observed individual relative accumulation of KIC/KMV, valine, leucine, and isoleucine (**Figure 4B-C**). We confirmed the significant accumulation of BCKA in CSF of patients with lymphoma LMD (n = 5) compared to CSF of patients without CNS involvement (n=8) using targeted mass spectrometry analysis with stable isotope-labeled standards for individual BCKA (**Figure 4D**). Combined levels of BCKA as high as 77.66 μM were observed in patients with lymphoma LMD. Interestingly, this effect was not unique to lymphoma and was also observed in CSF of patients with breast cancer (n = 12) and melanoma (n = 12) LMD (**Figure 4E**). These data highlight a dramatic accumulation of BCKA associated with LMD from multiple tumor types. The toxic effects of BCKA on neurons, glial cells, and astrocytes have been demonstrated previously^60–62^. We next wondered if the identified BCKA levels in CSF of LMD would directly affect the viability and function of neurons and meningeal fibroblasts in the leptomeningeal environment. We noted a significant decline in neuronal metabolic activity and viability after exposure to BCKA in complete neuronal media (**Figure 4F, Supplemental Figure 7**). Reflecting the leptomeningeal microenvironment, we exposed primary neurons and meningeal cells to different concentrations of BCKA in the context of CSF that mimics the physiologic conditions (ion, protein, chemical composition, see *Methods*). In this context, we observed a decline in neuronal and meningeal metabolic activity with exposure to BCKA as low as 25μM (**Figure 4G-H**).

**Figure 4.**
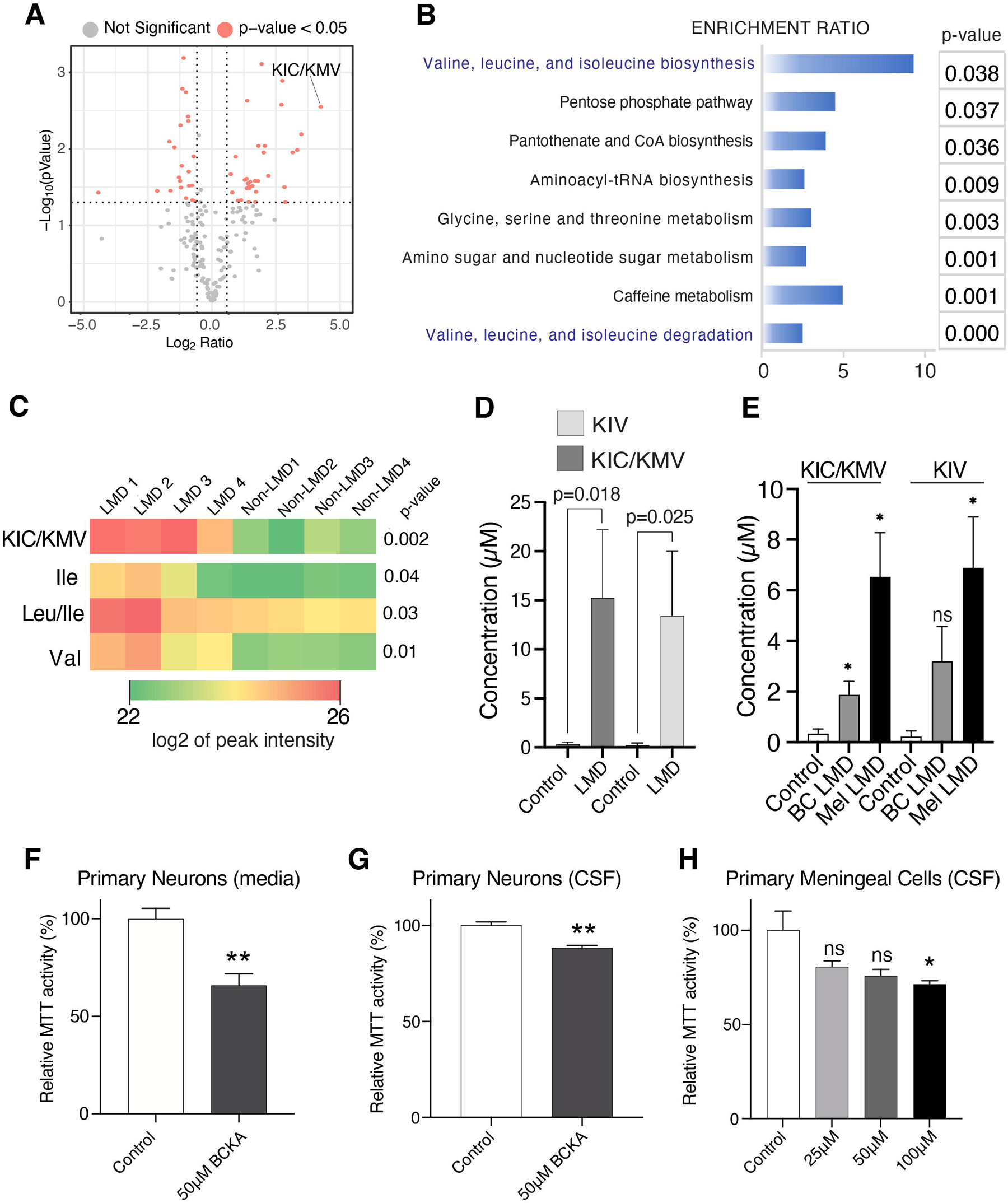
The metabolic environment of LMD is characterized by BCKA accumulation. A. Volcano plots show differentially abundant metabolites in the CSF of lymphoma patients with LMD compared to non-LMD controls. α-keto-isocaproic/ a-keto-β methyl valeric (KIC/KMV) isomers show the highest upregulated fold change. B. Pathway enrichment analysis of the differentially abundant metabolites using MetaboAnalyst 5.0 bioinformatics tool. C. Heatmap for the individual relative expression of branched-chain amino acids and branched chain-keto acids in lymphoma-LMD and non-LMD measured by mass spectrometry. D. The absolute concentration of branched-chain ketoacids in lymphoma-LMD vs.non-LMD measured by mass spectrometry. E. The absolute concentration of branched-chain ketoacids in breast cancer LMD and melanoma-LMD vs. non-LMD measured by mass spectrometry. F. MTT assay measuring metabolic activity of primary murine neurons in response to BCKA exposure in neuronal culture media for seven days. G. MTT assay measuring metabolic activity of primary murine neurons in response to BCKA exposure in physiological CSF for seven days. H. MTT assay measuring metabolic activity of primary human meningeal cells in response to BCKA exposure in physiological CSF for 72 hours. Data represent mean ± SEM (panels D, E, F, G, and H). Statistical significance was assessed using Welch’s t-test (C) and Student’s t-tests (panels D, E, F, G, and H). Significance is denoted as *p < 0.05, **p < 0.01, ***p < 0.001, ****p < 0.0001, ns = not significant.

### Branched-chain keto acids inhibit T-cell viability and function

BCKA accumulation results in a constellation of symptoms for patients with Maple Syrup Urine Disease, including immune dysfunction ^63^, but the direct mechanisms are poorly understood. We decided to directly test whether BCKA accumulation may affect T cell viability and function. Treating CD4 and CD8 T-cells isolated from human donors with concentrations of 50 μM to 10 mM BCKA at a physiological ratio (1:1.6:2.2 for KMV:KIV:KIC) in regular T cell media reduced T cell proliferation at higher BCKA doses following stimulation (**Figure 5A, B**). However, treating T cells with BCKA in the context of physiological CSF blunted T cell proliferation even at 50 μM doses (**Figure 5C, D**). BCKA treatment of T cells in physiological CSF severely reduced the secretion of pro-inflammatory effector cytokines, including interferon-γ, TNF-α, granzyme B, and interleukin-2 (**Figure 5E**). Individual KIC and combined BCKA reduced the viability of healthy donor T cells even at 25-50 μM doses after a 48-hour exposure in physiological CSF (**Figure 5F**). We also observed a slight increase in the expression of LAG3 checkpoint protein. However, this increase was not statistically significant (**Supplemental Figure 8**). This change was consistent with the elevated expression of LAG3 observed in CD8 T cell cluster 4, a major cluster common to all lymphoma LMD patients (**Figure 2A-B**). Similar effects on viability and cytokine secretions were observed in CD19-targeting CAR T cells (**Figure 5G-H**). On the other hand, BCKA treatment did not alter the viability of established lymphoma tumor cell lines, with the majority of the cell lines showing no effects on viability (**Supplemental Figure 9A-B**). These data suggest that accumulation of BCKA within the LMD tumor microenvironment induces an immune-suppressive microenvironment favoring tumor growth and survival.

**Figure 5.**
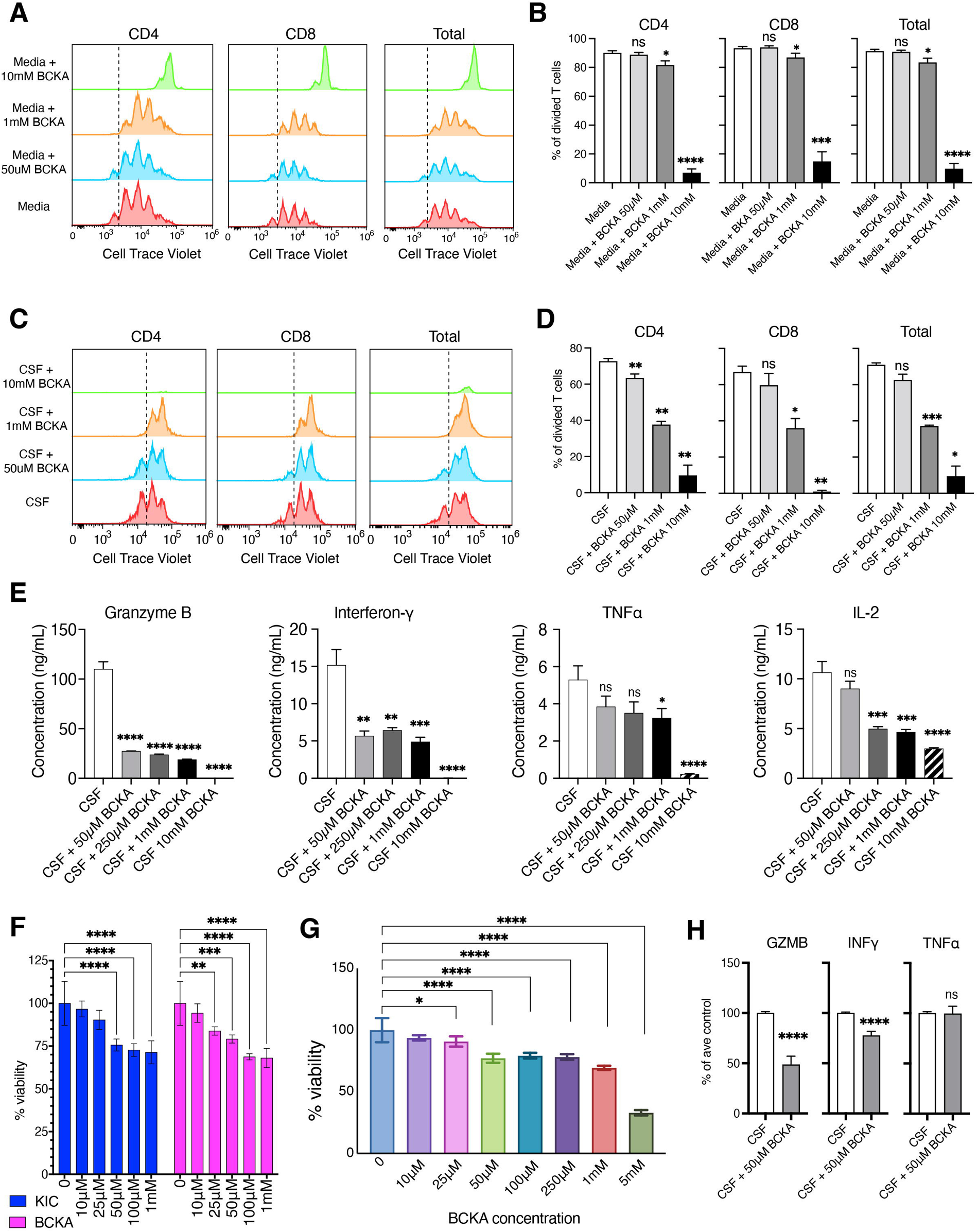
Impact of BCKA on T-cell viability and function. A. Representative proliferation plots of cell-trace violet labeled T-cells after OKT3, CD28, and IL-2 stimulation in full media with different concentrations of BCKAs or PBS (control). The experiment was repeated using three different PBMC donors. B. The percentage of T cell division was calculated from panel A using FlowJo. C. Representative proliferation plots of cell-trace violet labeled T-cells after OKT3, CD28, and IL-2 stimulation in physiological CSF with different concentrations of BCKAs or PBS (control). The experiment was repeated using three individual PBMC donors. D. The percentage of T-cell division was calculated from panel C using FlowJo. E. ELLA assays measuring the abundance of pro-inflammatory cytokines secreted from T-cells cultured in physiological CSF. This experiment was repeated with technical and biological triplicates using three different PBMC donors. F. Viability of human T-cells measured by Calcein AM staining in response to different concentrations of α-ketoisocaproic acid (KIC) or the three branched-chain keto acids (BCKA, at physiological ratio) in physiological CSF for 48 hours. This experiment was repeated twice using independent PBMC donors. G. Viability of human anti-CD19 CART-cells measured by Calcein AM staining in response to different concentrations of branched-chain keto acids (BCKA) in physiological CSF for 72 hours. This experiment was repeated using two independent PBMC donors. H. ELLA assays measuring the abundance of pro-inflammatory cytokines secreted from human anti-CD19 CART-cells cultured in physiological CSF. This experiment was repeated with technical triplicates and biological duplicates using two different PBMC donors. Data represent mean ± SD (panels B, D, E, F, G, and H). Statistical significance was assessed using Student’s t-test (B, D, E, F, G, H). *p < 0.05, **p < 0.01, ***p < 0.001, ****p < 0.0001, ns = not significant.

### Neurological decline in lymphoma LMD mouse model

The major debilitating symptoms of LMD from any tumor type result from neurological dysfunction ^64,65^. To quantify if our animal models of LMD display neurocognitive decline analogous to human disease, we employed neurological assessments commonly utilized in the study of neurodegenerative diseases, including tail suspension test (hind limb splay), grip strength test, motor function test, grooming, and kyphosis (**Supplementary Table 14**) ^66–68^. We have observed a significant reduction in the ability of animals with lymphoma LMD to perform a normal hind limb splay when suspended by the tail (**Figure 6A**). Likewise, their grip strength became significantly weaker with disease progression, with a profound loss in neuro-motor coordination, which can manifest as ataxia or paralysis (**Figure 6B**). In addition, malignant manifestations of the spinal cord can have profound effects on the structural stability of the spine and lead to kyphosis (excessive curve of the spine) in patients with LMD ^69,70^. We observed prominent kyphosis in the animals with LMD (**Figure 6C**). Even the skulls of animals with LMD showed a significant deformity in shape (**Supplemental Figure 10A**). Most mice with LMD reach the endpoint based on neurological complications three to four weeks after tumor engraftment (**Figure 6D**). During necropsy, significant degradation of the pia membrane was noted in the LMD animals (**Figure 6E**). We were unable to locate the transparent pia membrane (denoted by green arrow) on these animals and noted extensive tumor deposits instead (denoted by blue arrow) (**Figure 6E**). This observation was consistent with a dramatic reduction of fibroblast-like cells in the LMD pia enumerated using scRNAseq (66% of all cells in healthy pia versus 3% of all cells in LMD, **Supplemental Figure 10B**). Previously, neurological degeneration was associated with a loss of cortical microtubule-associated protein 2 (MAP2) reactivity and gliosis marked by an increase in GFAP expression in schizophrenia, Parkinson’s disease, Alzheimer’s disease, and other neurodegenerative processes ^71,72^. We also observed a loss of MAP2 reactivity in the cortex of animals with LMD (**Figure 6F**). This loss was coupled with increased GFAP staining of brain tissues, with increasing enhancement closest to the meningeal surfaces of the brain (**Supplemental Figure 10C**). Analysis of the digested pia tissues confirms an accumulation of BCKA in animals with LMD compared to PBS-injected controls (**Figure 6G**).

**Figure 6.**
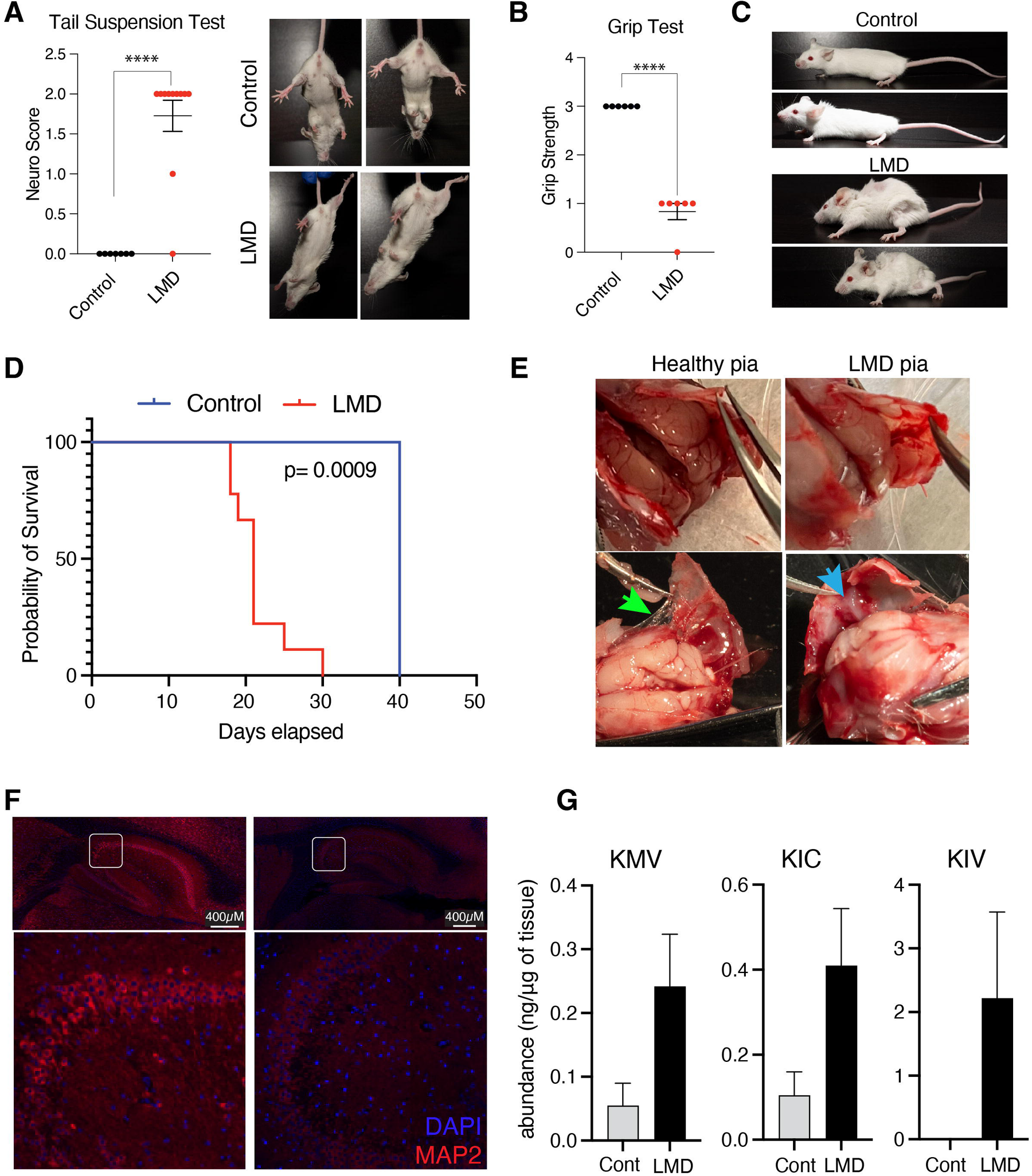
Neurodegenerative pathology of LMD. A. Tail suspension scoring for control PBS-injected mice vs. A20-injected lymphoma LMD mice. B. Grip strength assessing fore limb function for control PBS injected mice vs. A20 injected lymphoma LMD mice. C. Manifestation of kyphosis and loss of grooming in control PBS-injected mice vs. A20 injected lymphoma LMD mice. D. Kaplan-Meier analysis for probability of survival in control PBS injected mice vs. A20 injected lymphoma LMD mice. The p-value was calculated using the Log-Rank (Mantel-Cox) test. E. Images show a healthy, transparent pia membrane in the healthy control animals (green arrow) and a lack of this membrane in the LMD animals. LMD animals show significant tumor deposits instead (blue arrow). F. Immunofluorescent images of microtubule-associated protein 2 (MAP2, red) and DAPI (blue) in the brain hippocampus of LMD-lymphoma model. The top images show low magnification (scale bar is 400 μm), and the bottom images show high magnification of outlined areas. The scale bar is 400 μ m. Staining was repeated on four sections from different mice. G. Quantification of BCKA levels in brains of control and LMD-lymphoma model. Levels are calculated as ng/μg tissue using mass spectrometry. Data represent mean ± SEM (panels A, B, and G). Statistical significance was assessed using the Student’s t-test. Statistical tests were two-sided. ****P < 0.0001, ns = not significant.

### BCKA-reducing therapy extends survival and reduces neurological deterioration in the lymphoma LMD model

Most patients with LMD deteriorate rapidly regardless of treatment, highlighting an urgent clinical need to comprehend the immune-suppressive LMD biology and develop improved treatment strategies for LMD^73^. Despite the effectiveness of CD19-targeting CAR T-cell therapy in treating chemo-refractory B-cell lymphoma^20–24^, the responses in LMD patients are short-lived^25^. Our *in-vitro* experiments demonstrate the immunosuppressive effects of BCKA on the effector function of T cells and CAR T cells. We next investigated whether BCKA-reducing therapy improved the efficacy of CAR T cell treatments in our LMD mouse model. Sodium phenylbutyrate (PBA) is an FDA-approved drug used to treat inborn metabolic diseases that result in the accumulation of BCKA and branched-chain amino acids, such as the maple syrup urine disease ^74–76^. Mice were injected with A20 (lymphoma cells) subcutaneously and intrathecally. We then treated the mice with either anti-CD19 CART therapy with or without PBA (**Figure 7A**). Although the expected slight decline in weight had been observed for all animals with LMD at endpoint, no significant treatment-related toxicity was observed in any treatment group, but PBA-treated animals showed the least disease-related weight loss (**Supplemental Figure 10E**). Notably, PBA as a monotherapy effectively extended both the overall survival and the progression-free survival (Neuroscore <2) compared to the untreated group (**Figure 7 B-C**) and delayed the onset of neurological symptoms (**Figure 7D**). Intriguingly, combining PBA with CART therapy substantially prolongs survival (**Figure 7 B-C**) compared to CAR T monotherapy. Mice treated with the combination demonstrated the best neurological function scores out of any treatment arm at endpoint, highlighting the value of using PBA to improve the quality of life with LMD even during eventual progression (**Figure 7D**, top). Consistent with the enhanced neurologic function in treated animals, PBA treatment blocked the loss of MAP2 reactivity in the cortex of animals with LMD, validating the effect of PBA preserving neuronal integrity in the LMD mouse model (**Figure 7E**).

**Figure 7.**
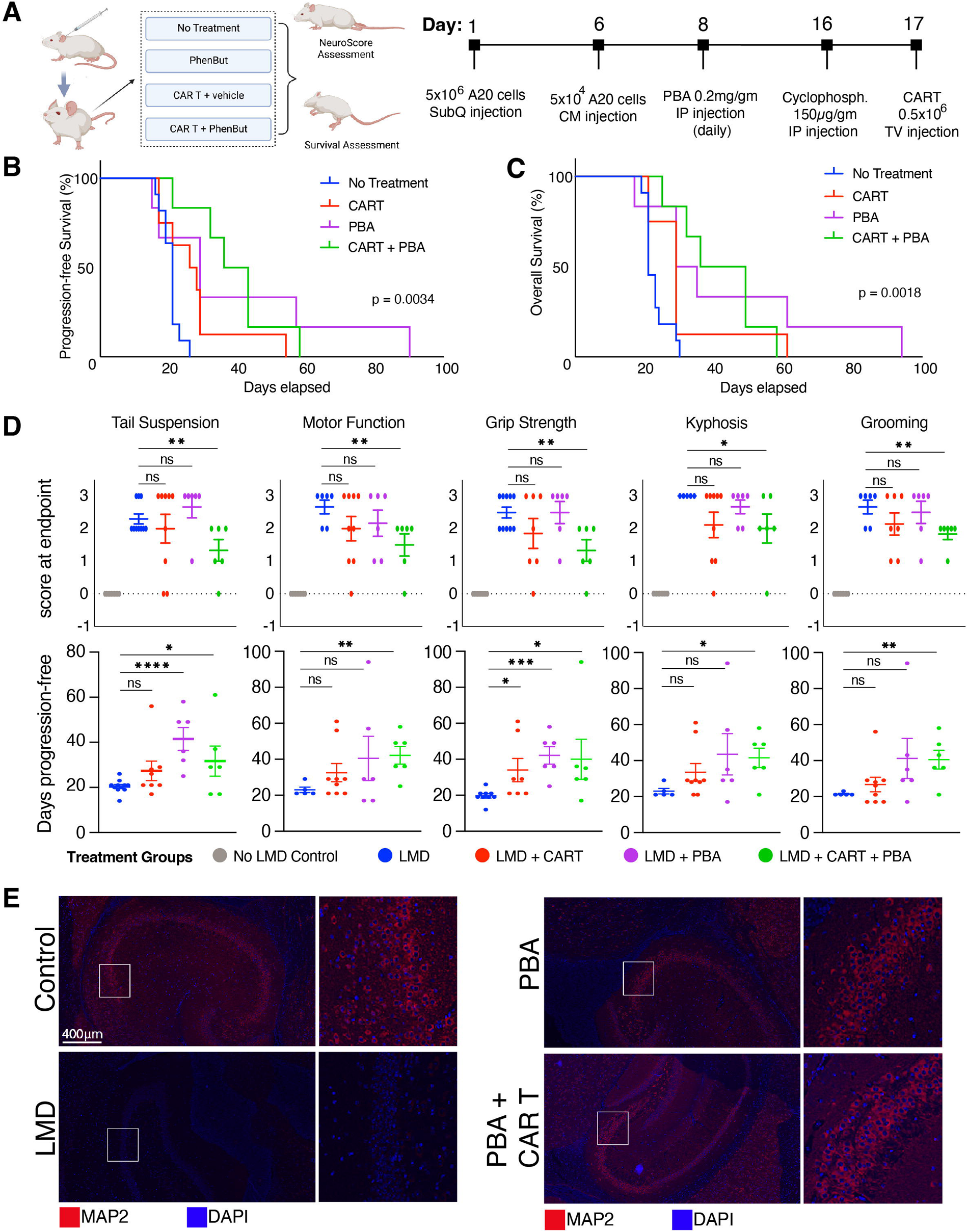
Sodium phenylbutyrate improves neurological functions and survival outcomes in lymphoma LMD mouse model. A. Schematic illustration for the *in vivo* experimental strategy showing the four LMD (A20 cell line injected) groups (left) and the timeline of tumor injections and the administered therapy (right). B. Kaplan Meier graph showing the progression-free survival for all mice. Progression was diagnosed in mice who scored a two or higher in any neurological assessment (motor function, tail suspension, kyphosis). The p-value was calculated using the Log-Rank (Mantel-Cox) test. C. Kaplan Meier graph showing the overall survival for all mice. The p-value was calculated using the Log-Rank (Mantel-Cox) test. D. Individual graphs showing the individual scores for each neurological assessment at endpoint (or last assessment before death), top. Individual graphs showing the number of days until mice displayed a score of 2+ for each assessment, paralysis or death, bottom. E. Immunofluorescent images of microtubule-associated protein 2 (MAP2, red) and DAPI (blue) in the brain hippocampus of the lymphoma LMD model. The left images show low magnification (scale bar is 400 μm), and the right images show high magnification of outlined areas. Staining was repeated on four sections from different mice. Data represent mean ± SE (D). Statistical significance was assessed using the Log-rank test (B, C) and Student t-test (D). *P < 0.05, **P < 0.01, ***P < 0.001, ****P < 0.0001, ns = not significant.

## DISCUSSION

LMD is a devastating complication with a dismal prognosis due to rapid neurological deterioration and with survival of untreated disease counted in mere weeks ^6^. Despite the small sample size as a limitation, we used single-cell transcriptomics and bulk multi-omics techniques to characterize the cellular and fluid compartments of the TME to gain novel insights into the biology of this rare but deadly disease. We uncovered a surprising accumulation of BCKA in the CSF tumor microenvironment. We then functionally validated the immunosuppressive and neurodegenerative role of BCKA accumulation in LMD using the *in vitro* and *in vivo* models.

Analyzing the transcriptional landscape of the CSF samples revealed a profound cellular enrichment in the CSF of patients with LMD, including tumor B-cells, T-cells, and myeloid cells, indicating direct tumor-immune interactions within the CSF space. The CSF of patients with short survival retain an immunosuppressive microenvironment involving the accumulation of tumor-promoting (alternatively activated) macrophages, a dysfunctional T cell landscape, and the absence of antigen-presenting dendritic cells. On the other hand, the CSF samples from the extraordinary survivor showed few macrophages, subsets of activated and proliferating T-cells, and multiple dendritic cells. Previously, tumor-promoting macrophage infiltration in the tumor microenvironment has been attributed to the progression of glioblastoma ^77^, lung cancer ^78^, and sarcoma ^79^. Interestingly, the accumulation of the tumor-promoting macrophages was not limited to lymphoma-LMD but was also confirmed in a previously published cohort of CSF from melanoma LMD patients ^26^. Additionally, mouse models of breast cancer, melanoma, and lymphoma LMD demonstrated a significantly higher macrophage accumulation within the leptomeningeal microenvironment than the extracranial disease, further validating the accumulation of macrophages in the LMD tumor microenvironment.

The fluid microenvironment of the TME likely governs the cellular landscape and the cell-cell interactions. The analysis of CSF from lymphoma LMD patients showed significant alterations in the abundance of ∼30% of lipids, proteins, and metabolites. Interestingly, multi-omic data analysis suggested a strong link between the fluid tumor microenvironment and the immune-suppressive and neurodegenerative processes at this unique tumor site. We observed an upregulation in lysophosphatidylcholines, which are directly correlated with neuron demyelination ^44,45^. On the other hand, LMD CSF showed a depletion in the membrane lipid class, phosphatidylethanolamines (PE). PE deficiency has been reported in CSF collected from idiopathic Parkinsonism and has also been correlated with lipid-induced ER stress and aging ^80,81^. Likewise, the proteomic analysis of LMD CSF revealed a depletion in numerous proteins (including GRIA4, ARSA, THY1, and TAC1) vital for proper neurological function. Focusing on proteins with significant alterations in Alzheimer’s, Parkinsonism, or multiple sclerosis, LMD CSF samples were significantly enriched with immunoglobulin G, transferrin, vitamin-D binding protein, PARK7, and DJ-1, while amyloid β, tachykinins, and NPTXR were significantly depleted. Additionally, proteomic analysis demonstrated the enrichment of proteins involved in complement cascade activation (C2, C5, FCN2, FCN3, CPN1, and MBL2), which have been previously reported with melanoma LMD ^82^. Another study has reported C3 as a contributor to tumor dissemination via blood-CSF barrier disruption in LMD with breast and lung cancer ^83^. These data strongly suggest a significant role of complement activation cascade in LMD.

The most profound discovery stemmed from the analysis of the fluid metabolomics, where we discovered a tremendous accumulation (∼45-60-fold average increase over control) of BCKAs in LMD. BCKAs are derived from the deamination of the branched-chain amino acids ^84^. Accumulation of BCKAs is linked to the neurotoxicity identified in Maple Syrup Urine disease (rare inborn error of metabolism disorder) ^84–88^. Although limited studies investigated the role of BCKAs in cancer ^89,90^, the elevation of BCKAs has never been reported in LMD, and the effects of these toxic metabolites on T-cell function have never been investigated. Our data reveals that BCKAs can significantly reduce both the viability, proliferation, and effector function of endogenous T cells and anti-CD19 CAR T cells. These results outline the contribution of BCKA accumulation to the immune-suppressed microenvironment of LMD.

The rapid and debilitating neurological symptoms are the leading cause of poor survival in LMD patients ^1,91^. Our LMD mouse model showed similar symptoms for neurological decline coupled with the loss of leptomeningeal integrity, neuronal dysfunction (loss of MAP2), and induction of reactive (GFAP+) astrocytes. Although some of these symptoms may be caused by hydrocephalus, we have observed a direct inhibitory effect of BCKAs on the meningeal and neuronal cells’ metabolic function. Although the toxic effects of BCKAs on the brain’s neuronal, glial cells, and astrocytes have been reported before, we demonstrate that these effects occur at much lower concentrations than previously tested (100-500x lower) ^63,88,92^. BCKA-lowering therapy, such as PBA, has been shown to improve neurological symptoms in MUSD patients ^74–76^. Thus, we examined whether PBA would alleviate some of the severe neurodegenerative processes and prolong the survival of animals with LMD. Intriguingly, PBA shows a profound improvement in the neurological functions and the survival of the mice, confirming the role of BCKAs in LMD pathogenesis. Furthermore, PBA treatment also enhanced the efficacy of anti-CD19 CAR T cell therapy, with the combination showing the highest median overall survival, highest median progression-free survival, and lowest manifestation in neurological symptoms. These data indicate the possible influence of BCKA accumulation in LMD on blunting the long-term efficacy of CAR T therapy.

Overall, we uncover the role of BCKAs as toxic metabolites inducing an immune suppressive and neurodegenerative tumor microenvironment in lymphoma LMD. We provide the first evidence that BCKA-reducing strategies have the potential to improve the quality of life, survival outcomes, and efficacy of CAR T cell therapies in patients with LMD.

## METHODS

### Patient specimens

This study was conducted by recognized ethical guidelines (e.g., Declaration of Helsinki, CIOMS, Belmont Report, U.S. Common Rule). Eight CSF specimens were procured in this study under protocols approved by the Institutional Review Board (MCC#19332) (**Table 1**). Four specimens were collected from lymphoma LMD patients, and four were from non-LMD patients. All CSF samples were obtained during routine clinical care (**Supplemental Figure 1**). Samples were placed on ice and processed immediately. CSF was spun down to separate the fluid and cellular compartments. The fluid was preserved in 500 μL aliquots at -20^0^C, and the cells were cryo-preserved as viable suspensions in 10% DMSO/FBS.

### Cell culture

All Cells were maintained at 37 °C in a 5% CO_2_ atmosphere.

#### Lymphoma cells

The murine lymphoma A20 cell line ^93^ was purchased from ATCC. OCI-LY3, Toledo, SUDHL4, and JEKO are human lymphoma cell lines. All lymphoma cell lines were cultured using RPMI-164 medium supplemented with 10% heat-inactivated fetal bovine serum (Sigma-Aldrich), 1mM sodium pyruvate (CORNING), 1x MEM Nonessential Amino Acid (CORNING), and 0.05 mM 2-mercaptoethanol (Sigma-Aldrich).

#### Primary T cells

Human T cells were isolated from peripheral blood mononuclear cells (PBMC) using EasySep Human T Cell Iso Kit (STEMCELL), while mouse T cells were isolated from Balb/C mouse spleen using EasySep Mouse T Cell Isolation Kit following the manufacturer’s instructions. Then, T-cells were cultured in 10% FBS-RPMI with 1% sodium pyruvate, 1% nonessential amino acids, and 1% Pen-Strep.

#### CD19–28z CART generation

T cells were stimulated with Dynabeads T-Activator CD3/CD28 (Gibco), 200IU/mL of IL-2 (PeproTch) in full media. The next day, in retronectin-coated 6-well plates, stimulated T cells were cultured with retroviral supernatant collected from Phoenix E (Mouse) or RD114 (Human) producer cells to transduce mouse or human T cells as previously described ^94^. Transduction efficiency was detected using flow cytometry as a percentage of mCherry^+^ or Ametrine**^+^** cells.

#### Primary neurons

Primary Rat Cortex Neurons (A36511, Gibco) were cultured in a poly-L-lysine-coated 96-well plate. The culture media consisted of Neurobasal Plus Medium containing 1X B27 (50X, Gibco), 200mM Glutamax, and 20mM HEPES.

#### Primary human meningeal cells

Primary human meningeal cells (ScienCell) were grown in Meningeal cell media containing 2% FBS, 1% growth supplements, and 1% antibiotics.

### Physiological cerebrospinal fluid (CSF)

To prepare CSF with physiological properties, synthetic CSF (TOCRIS) was supplemented with 1% fetal bovine serum, 60mg/dL D-Glucose (Sigma-Aldrich), 1x MEM Nonessential Amino Acids (CORNING), and 15mM HEPES (LONZA). CSF was then filtered with a 0.22 μM filter and was used within one day of preparation.

### Branched-chain keto acids

3-methyl-2-oxovaleric acid sodium salt (KMV) (Toronto Research Chemical), 4-methyl-2-oxovaleric acid (KIC) (Sigma), and Sodium 3-methyl -2-oxobutyrate (KIV) (Sigma) were used to prepare branched-chain ketoacid (BCKA) solutions (in PBS) in the ratio 1:2.2:1.6 to recapitulate the physiological concentrations^95^.

### T cell proliferation assay

Forty-eight well plates were coated with OKT3 (5 μg/mL), then 2 x 10^5^ fresh PBMC, prelabeled with cell trace violet (CTV), were cultured in 500 μL of full media containing CD28.2 (2 μg/mL) and IL2 (200 IU/L) for 24 hours. The next day, 400 μL of the media was carefully removed, and 350 μL of either full media or physiological CSF was added with 50 μL BCKA (10X the required concentration) or PBS vehicle control.

### Viability test

T cells: In black 96 well plates, pre-coated with OKT3 (5 μg/mL), 3 x 10^4^ T-cells were cultured in full media and IL-2 for one day, and then the media was changed to either fresh full media or CSF with BCKA or PBS vehicle control. **Lymphoma cells and CAR T**. Human lymphoma cells (3x10^4^) were plated in black 96-well plates in 40 μL full media. Then, 140 μL of physiological CSF and 20 μL of 10X BCKAs (PBS for controls) were added to each well. After 48 hours, Calcein Am (Invitrogen) was added for one hour at 37 °C, and the fluorescence intensity was measured at 485nm excitation and 520 nm emission wavelengths.

The viability of lymphoma cells was validated with propidium iodide staining. In 24 well plates, four human lymphoma cell lines (Toledo, JEKO, OCI-LY3, SUDHL4) were cultured at density 2. 5x 10^5^ in 100 μL full media with 350 μL CSF and 50 μL 10X BCKA. After 48 hours, cells were collected and stained with propidium iodide and analyzed by flow cytometry.

#### MTT assays

For neurons, a 96-well plate was coated with poly-L-lysine, and then neurons were cultured in neurobasal media or physiological CSF. Then, 50 μM BCKA or PBS vehicle control was added. On the 3^rd^ day, the media was changed with fresh media with BCKA or PBS vehicle control. On the 7^th^ day, 20 μL of MTT (5 mg/mL) solution was added to each well and incubated at 37 °C for four hours, then 200 μL of DMSO was added to dissolve formazan crystals. Absorbance was read at 550 nm. For meningeal cells, they were cultured in a 96 well-plate in full media for 72 hours. Then, the same protocol for the MTT assay was performed.

### ELLA multiplex cytokine assay

Forty-eight well plates were coated with OKT3 (5 μg/mL), then 2 x 10^5^ fresh PBMC were cultured in the presence of CD28 (2 μg/mL) and IL2 (200 IU/L) for 24 hours, and 500 μL of full media. The next day, 400 μL of the media was carefully removed, and 350 μL of either full media or physiological CSF was added with 50 μL BCKA (10X the required concentration) or PBS vehicle control. On the fifth day of culturing, 300 μL of the supernatant was collected to measure Granzyme B, Interferon, TNF, and IL-2 using Ella multiplex assay plate (Bio-techne). The cells were stained using the following anti-human antibodies: CD4 (BUV395), CD8 (FITC), PD-1(PE), and LAG3 (BV421). Live/Dead (NIR) stain was used for viability. Samples were analyzed on an LSR II flow cytometer (Beckman Coulter), and all data were analyzed using FlowJo.

### Immunohistochemistry

Mouse brains were collected immediately after euthanasia. The two brain hemispheres were separated using a scalpel. One hemisphere was fixed in 10% neutral buffered formalin for ∼ 24 hours; then, brains were transferred to aqueous 70% ethanol and kept at 4 °C until sectioning. Mouse brains were embedded in paraffin blocks and sectioned (4-micron thickness) at Moffitt’s Tissue Core facility. Brain sections were rehydrated, then blocked with goat serum (Gibco) for one hour, followed by incubation with either rabbit anti-GFAP (1:1000) or rabbit anti-MAP2 (1:1000) antibody overnight at 4 °C. Alexa Fluor 680 goat IgG (Invitrogen) was used as the secondary antibody. Slides were mounted using SlowFade Diamond mounting media containing DAPI and imaged on the Akoya slide scanning microscope. Images were visualized using Phenochart 1.2.0 software.

#### Animal studies

All mouse work was conducted in accordance with recognized ethical guidelines and IACUC approval. Female BALB/c mice at 12 weeks of age were used to establish a murine model of lymphoma LMD. The murine lymphoma LMD group was injected with A20 cells in both the cisterna manga (5 x 10^4^ cells in 5 μL PBS) and the left side lymph node of the leg flank (concentration 5 x 10^5^ cells in 30 μL PBS) or subcutaneously (5 x 10^6^ cells in 100 μL PBS). The control group was injected with only PBS in both the cisterna magna (5 μL) and the left side lymph node of the leg flank (30 μL) or subcutaneously (100μL PBS). The extracranial metastasis group was injected intravenously in the tail vein with either A20 cells (concentration 5 x 10^4^ cells in 100 μL PBS) or 100 μL of PBS as a control. Mice were anesthetized during the procedure and received analgesics for three days to alleviate any pain from the surgery. Mice were weighed twice a week and evaluated using neurological function assessments. For the neurological evaluation, we used a group of tests previously described in neurodegenerative diseases ^66–68^. These tests are 1) hind limb splay, which assesses the ability of the mouse to control the whole-body muscles and perform a hind leg splay during a tail suspension, 2) motor function, which asses the neuro-motor coordination and function, 3) Grooming is a behavioral test for the overall health status of the mouse, 4) Gripping of the wire lid, which is an indicator of the strength of the fore limb, 5) Kyphosis is used to observe spine curvature or deformity and has been used to evaluate neurodegenerative diseases in mouse models. Anti-CD19 CAR T-treated mice had lymphodepletion using cyclophosphamide (150 μg/gm body weight, IP injection) 24 hours before CAR T injection. CAR T transfection efficiency was 60% (confirmed by mCherry-positive cells, **Supplemental Figure 10F**), and mice received ∼0.5 x 10^6^ CART via tail vein injection. Sodium Phenyl Butyrate (PBA)-treated mice received daily IP injection (0.2 mg/gm body weight in PBS). Mice were euthanized by cervical dislocation when they lost more than 20% of their original weight or showed severe neurological decline such as paralysis or ataxia.

#### Single-cell RNA-seq

Cryopreserved cells were carefully recovered from freeze and strained to obtain a single-cell suspension. Cell number and viability were assessed using Countess II FL automated cell counter. Cells were resuspended in cold 0.2% BSA/PBS at an optimum concentration (500 cells/µl). The cell suspension was directly loaded for scRNA-seq library preparation using the Chromium Single Cell Reagent Kit following the manufacturer’s protocol (10x Genomics, USA) as previously published ^26^. The cDNA libraries were prepared using the Single-Cell 3′ Library Prep Kit (10X Chromium). The resulting libraries were sequenced on the Illumina NextSeq 500 instrument using v2.5 flow cells. Approximately 80,000 to 1,000,000 mean sequencing reads per cell were generated. scRNA[seq sequencing data were demultiplexed, aligned, and quantified using the 10X Genomics Cell Ranger Single[Cell Software Suite against the human reference genome. Quality control and cell type analysis were performed as previously described ^26^. We computed the Ligand-Receptor interactions between immune and tumor cells via CellPhone DB 3.0.1 (Python-based computational analysis tool)^96^. The resulting datasets from patients with long and short survival were uploaded on an interactive R/Shiny-based platform called InterCellar^97^. InterCellar allows for the comparison of both datasets and provides unique cell-cell interactions.

#### Proteomic, Metabolomic, and Lipidomic analysis

Cell-free cerebral spinal fluid (CSF) samples were sent to the Proteomics & Metabolomics Core at Moffitt Cancer Center to perform Proteomic, Metabolomic, and Lipidomic analyses.

LC-MS grade solvents and additives, including water, methanol, acetonitrile, and formic acid, were purchased from Burdick and Jackson, Honeywell, Muskegon, MI (sourced through VWR), and Thermo Scientific. Neat standards 4-methyl-2-oxovaleric acid, 3-methyl-2-oxovaleric acid, 3-methyl-2-oxobutanoic acid, and Glutamic acid were purchased from Millipore Sigma. Stable isotope-labeled standards (SIS) were obtained from Millipore Sigma and Toronto Research Chemicals; it includes 3-Methyl-2-oxobutanoic acid (^13^C_2_, 98%; D, 99%), 4-Methyl-2-Oxovaleric acid (D3, 98%), Glutamic acid (^13^C_5,_ 98%; ^15^N, 98%) and 3-Methyl-2-Oxovaleric acid (D8, 98%).

### Untargeted proteomic analysis

Proteomic analysis was performed as previously described ^98^. Briefly, the CSF (cell-free) supernatant was concentrated using an Amicon Ultra membrane filter (Millipore), followed by depletion of the top 12 abundant protein spin columns (Pierce). The Flow-through underwent reduction, alkylation, and digestion using Trypsin. Tryptic peptides (5 µg) were then labeled with TMT-11plex reagents (Thermo), following the manufacturer’s protocol. Label incorporation was verified to be >95% by LC-MS/MS and spectral counting of labeled versus unlabeled peptides (Protein Discoverer, Thermo). The samples were then quenched with aqueous 5% hydroxylamine, pooled, and lyophilized. The TMT channel layouts were: 127N-LMD_1, 128N-LMD_2, 129N-LMD_3, 130N-LMD_4, 127C-Normal_1, 128C-Normal_2, 129C-

Normal_3, 130C-Normal_4. Pooled TMT labeled peptides were fractioned using basic pH reversed-phase HPLC fractionation, and 24 concatenated fractions were collected using the following method. After lyophilization, TMT-labeled peptides were redissolved in 250 μL of aqueous 20 mM ammonium formate (pH 10.0) for high pH reversed-phase liquid chromatography (bRPLC) separation, performed on an XBridge 2.1 mm ID x 50 mm long column packed with BEH C18 resin, 3.5 µm particle size, 130 Å pore size (Waters). bRPLC Solvent A was aqueous 2% acetonitrile (ACN) with 5 mM ammonium formate, pH 10.0. Peptides were eluted using the following gradient program: 1% bRPLC B (aqueous 90% acetonitrile with 5 mM ammonium formate, pH 10.0) for 9 minutes, 1% – 9% B in 4 minutes, 9%-32% B in 25 minutes, 32%-55% B in 40 minutes, 55%-90% B in 3 minutes and 90% B held for 10 minutes, 90% to 1% B in 1 minute, and followed by re-equilibration at 1% B for 5 minutes. The flow rate was 0.2 ml/min. Vacuum centrifugation (Speedvac, Thermo) was used to dry the samples prior to redissolving in LC-MS/MS solvent A. A nanoflow ultra-high-performance liquid chromatograph and nanoelectrospray orbitrap mass spectrometer (RSLCnano and Q Exactive Plus, Thermo) were used for LC-MS/MS. The sample was loaded onto a pre-column (C18 PepMap100, 100 µm ID x 2 cm length packed with C18 reversed-phase resin, 5 µm particle size, 100 Å pore size) and washed for 8 minutes with aqueous 2% acetonitrile containing 0.1% formic acid. Trapped peptides were eluted onto the analytical column (C18 PepMap100, 75 µm ID x 25 cm length, 2 µm particle size, 100 Å pore size, Thermo). A 120-minute gradient was programmed as 95% solvent A (aqueous 2% acetonitrile + 0.1% formic acid) for 8 minutes, solvent B (aqueous 90% acetonitrile + 0.1% formic acid) from 3% to 30% in 80 minutes, from 30% to 38.5% in 10 minutes, then solvent B from 50% to 90% B in 5 minutes and held at 90% for 3 minutes, followed by solvent B from 90% to 5% in 2 minutes and re-equilibration for 15 minutes using a flow rate of 300 nl/min. Spray voltage was 1900 V. Capillary temperature was 250 °C. The top 16 tandem mass spectra were collected in a data-dependent manner. Settings for MS acquisition were 70,000 resolution, 3E6 AGC target, 200 ms Max IT, and recording m/z 440-2000. The settings for tandem mass spectrometry data acquisition were 35,000 resolution, 1E5 AGC target, 110 ms Max IT, isolation window 0.8 with 0.2 offset, fixed first mass at m/z 100, and 2 normalized collision energy (NCE) values of 24 and 29.

### Data Analysis

MaxQuant (version 1.6.14.0)^99^ was used to identify peptides using the UniProt human database (downloaded in June 2020) and quantify the TMT reporter ion intensities. Up to 2 missed trypsin cleavages were allowed. The mass tolerance was 20 ppm for the first search and 4.5 ppm for the main search. Reporter ion mass tolerance was set to 0.003 Da. Carbamidomethyl cysteine was set as a fixed modification. Methionine oxidation was set as a variable modification. Both peptide spectral match (PSM) and protein false discovery rate (FDR) were set at 0.01 (or 1%). The match between runs feature was activated to carry identifications across samples.

### Untargeted metabolomic analysis

100 µL of each CSF sample was spiked with 5 µL of a mixture of stable isotope-labeled standards (Cambridge Isotope Labs, Tewksbury, MA). An aliquot of pre-chilled methanol was added to each sample for a final composition of 80% methanol to extract metabolites and precipitate proteins. The samples were vortexed and incubated at -80 °C for 30 minutes. Proteins were removed from samples by centrifugation at 18,000 x g for 10 minutes at 4 °C. The supernatant containing the metabolites was lyophilized and then resuspended in 100 µL of 80% methanol. A 10 µL aliquot of each sample was used to make a pooled sample. Liquid chromatography-high resolution mass spectrometry (LC-HRMS) was performed using a UHPLC interfaced with Q Exactive HF. An aliquot (5 µL) of each sample was loaded onto a SeQuant ZIC-pHILIC guard column (4.6 mm inner diameter/ID x 20 mm length, 5 µm particle size) connected to a SeQuant ZIC-pHILIC column (4.6 mm ID x 150 mm length, 5 µm particle size) (MilliporeSigma, Burlington, MA), and maintained at 30 *°C*. For LC-MS analysis, mobile phase A contained aqueous 10 mM ammonium carbonate plus 0.05% ammonium hydroxide, and mobile phase B contained 100% acetonitrile. A linear gradient was programmed from 80 to 20% B over 13 minutes at a flow rate of 0.250 mL/min and then maintained at 20% B for 2 minutes, followed by re-equilibration over 5 minutes at a flow rate of 0.250 mL/min, for a total run time of 20 minutes for each experiment. The Q Exactive HF mass spectrometer was operated in positive and negative mode separately using a scan range from m/z 60 to m/z 900. LC-MS data files were converted to mzXML files using ProteoWizard and analyzed using MZmine 2.38 [1].

### Untargeted lipidomic analysis

A 200 µL aliquot of each CSF sample was spiked with 5 µL of SPLASH Lipidomix standard (Avanti Polar Lipids) and extracted with 600 µL of pre-chilled isopropanol. The samples were vortexed and incubated at -80 °C for 20 minutes. The protein was pelleted by centrifugation at 13,800 x g for 20 minutes at 4 °C, and its concentration was calculated using Bradford assays to estimate total protein content (Pierce™ Coomassie Protein Assay Kit). The supernatant containing the lipids was lyophilized and then resuspended in 100 µL of 100% methanol. The analysis was performed using LC-MS/MS with a Vanquish LC (Thermo, San Jose, CA) interfaced with a Q Exactive HF mass spectrometer (Thermo, San Jose, CA). Chromatographic separation was conducted on Brownlee SPP C18 column (2.1 mm x 75mm, 2.7 µm particle size, Perkin Elmer, Waltham, MA) using mobile phase A containing 100% water with 0.1% formic acid and 1% of 1M ammonium acetate and mobile phase B containing 1:1 acetonitrile: isopropanol with 0.1% formic acid and 1% of 1M ammonium acetate. The gradient was programmed as follows: 0-2 min 35% B, 2-8 min from 35% B to 80% B, 8-22 min from 80% B to 99% B, 22-36 min 99% B, 36.1-40 min from 99% to 35% B at a flow rate of 0.400 mL/min. Full MS and tandem mass spectrometry using data-dependent acquisition (top-10 method) were used in both positive and negative modes separately. Lipid data was analyzed using LipidSearch 4.2 (Thermo Fisher Scientific) and normalized by the amount of protein in the sample.

#### HPLC for branched-chain ketoacids

Sample preparations were carried out on ice. An aliquot (various amounts based on different amount starting materials) of the SIS mixture was added into each sample (either CSF or pulverized tissue). A pre-cooled 80% MeOH extraction solvent (kept in the -80 °C freezer at least one hour prior to extraction) was added to the sample for protein precipitation. After the addition of the extraction solvent, the samples were vortexed and then incubated for 30 minutes in a -80 °C freezer to increase metabolite extraction. After that, the samples were centrifuged at 18,800 × g (Microfuge 22R, Beckman Coulter) at 0 °C for 10 minutes. Then, the supernatant was transferred to a new microcentrifuge tube. For tissue samples, the protein pellet was resolubilized using aqueous 20 mM HEPES with 8 M urea for Bradford assays to measure the protein concentration. Dried Keto Acids and Glutamic acid were redissolved in aqueous 80% MeOH.

Ultra-high performance liquid chromatography-high resolution mass spectrometry (UHPLC-HRMS) was performed using a Vanquish UHPLC interfaced with a Q Exactive FOCUS quadrupole-orbital ion trap mass spectrometer (Thermo, San Jose, CA). Chromatographic separation was performed using ACE Excel 1.7 SuperC18 LC column (2.1 mm ID x 100 mm length, 1.7 µm particle size, Mac-mod Analytical, Chadds Ford, PA). Mobile phase A was aqueous 0.1% formic acid, and mobile phase B was 100% acetonitrile. The gradient program included the following steps: start at 5% B and stay for 1 minute, a linear gradient from 5 to 60% B over 5 minutes and then to 80% B within 0.5min, stay at 80% B for 1.5 minutes, return to 5% B in 0.1 minutes, and re-equilibration for 0.9 minutes for a total run time of 8 minutes. The flow rate was set to 0.400 mL/min. The autosampler was cooled to 5 °C, and the column temperature was set to 60 °C. The sample injection volume was 2 µL. Full MS was performed in negative mode, detecting ions from m/z 100 to m/z 200. Xcalibur, version 4.5, was used to identify and quantify metabolites by matching by m/z and RT.

#### Bioinformatic analysis

All of the identified features of proteomic (n = 1,410), metabolic (n = 193), and lipidomic (n = 160) datasets and their relative MS abundance were analyzed. The statistical analysis for our data was conducted as follows: 1) proteins, metabolites, and lipids intensities were Log_2_ transformed, 2) average Log_2_ abundances for LML and non-LML groups were calculated, 3) Log_2_ ratios were used to calculate the fold change of LML to non-LML, 4) Multiple t-tests were used for statistical comparison and false discovery rate (FDR <= 0.05) was used to determine significance. Principal component analysis (PCA) was used to visualize the data clustering from different patients. The significantly expressed features in the LMD group versus the non-LMD group were visualized using a heatmap and volcano plot. Ingenuity Pathway Analysis was used for pathway enrichment of proteins and metabolites between LMD and non-LMD samples with at least +/-1.5 fold change and p-value <= 0.05. ‘The Joint-Pathway Analysis’ module from MetaboAnalyst 5.0 (https://www.metaboanalyst.ca) was used for pathway enrichment of proteins and metabolites between LMD and non-LMD samples with at least +/-1.5 fold change and p-value <= 0.05. Results were downloaded and visualized in R (4.0.4) with RStudio (2023.06.1). The X-axis shows the pathway impact score (normalized topology measures of proteins and metabolites in each pathway), and the y-axis shows -log10(pathway p-value^100^.

## ACKNOWLEDGEMENTS

We thank the patients and their families for their essential and generous contributions to this study. The work in the Smalley lab is generously supported by the Research Scholar Grant from the American Cancer Society (RSG-23-1040487-01-MM), Tara Miller Melanoma Foundation Young Investigator Award from the Melanoma Research Alliance (932727), an R00 from the National Institutes of Health (R00 CA226679), and an R21 from the National Institutes of Health (R21 CA274060). MDJ is supported by the Mark Foundation, the Bankhead-Coley Cancer Research Program, and a Florida Academic Cancer Center Alliance grant. PAF has research support from CDMRP research grant, Department of Defense, Moffitt Cancer Center Center of Excellence Celgene project, NIH/NCI grant, Pfizer, State of Florida Bankhead Coley, NIH/NCI 1R21 grant, Biocept, Roswell Park Cancer Institute. This work has been supported in part by the Molecular Genomics Shared Resource, Proteomics and Metabolomics Shared Resource, and the Biostatistics and Bioinformatics Shared Resource at the H. Lee Moffitt Cancer Center & Research Institute, an NCI designated Comprehensive Cancer Center (P30-CA076292).

