## Supplemental Figures 1-10 for "Branched-chain keto acids promote an immune-suppressive and neurodegenerative microenvironment in leptomeningeal disease"

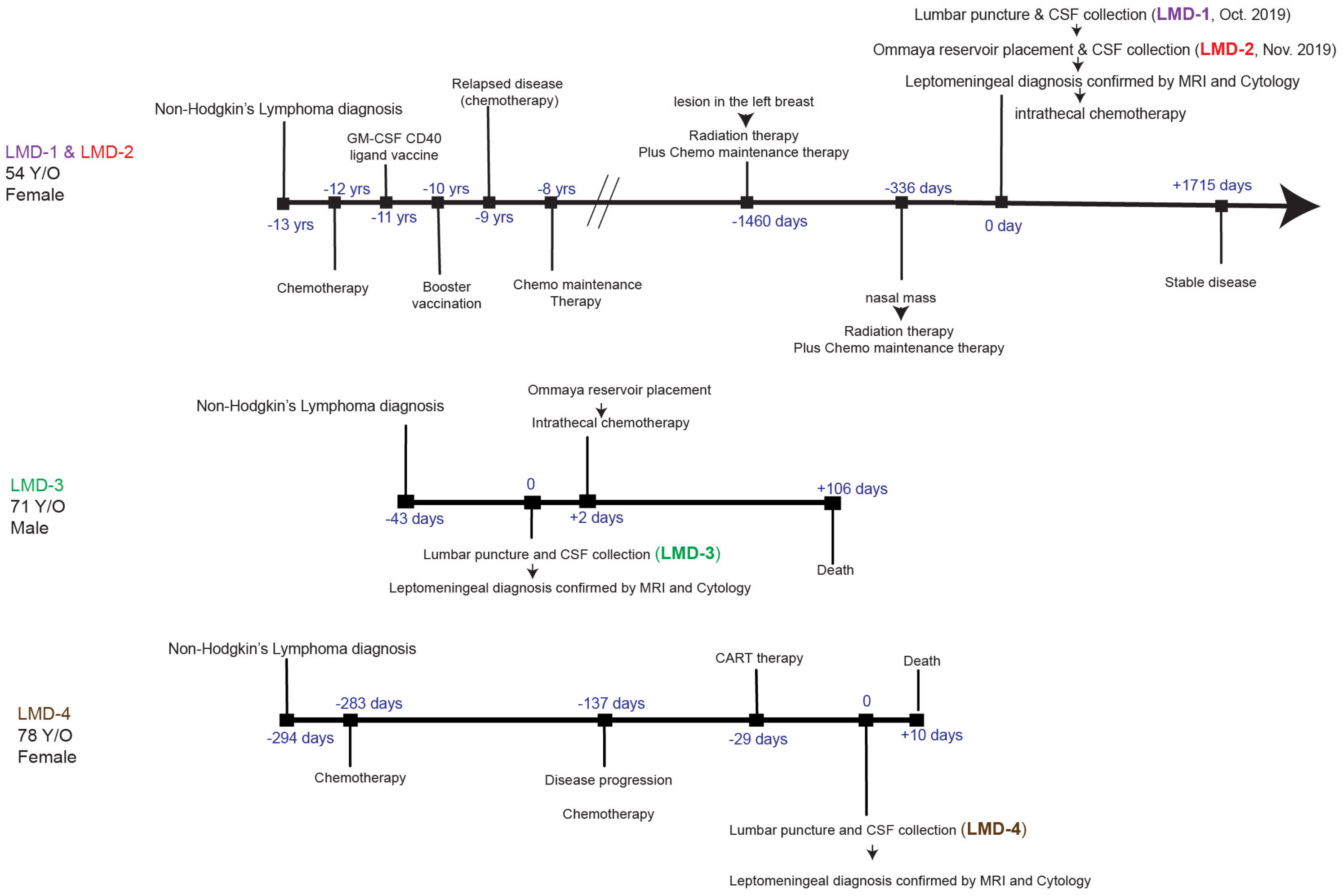


**Supplemental Figure 1.** Patient timelines showing relevant clinical milestones relative to CSF sample collection.

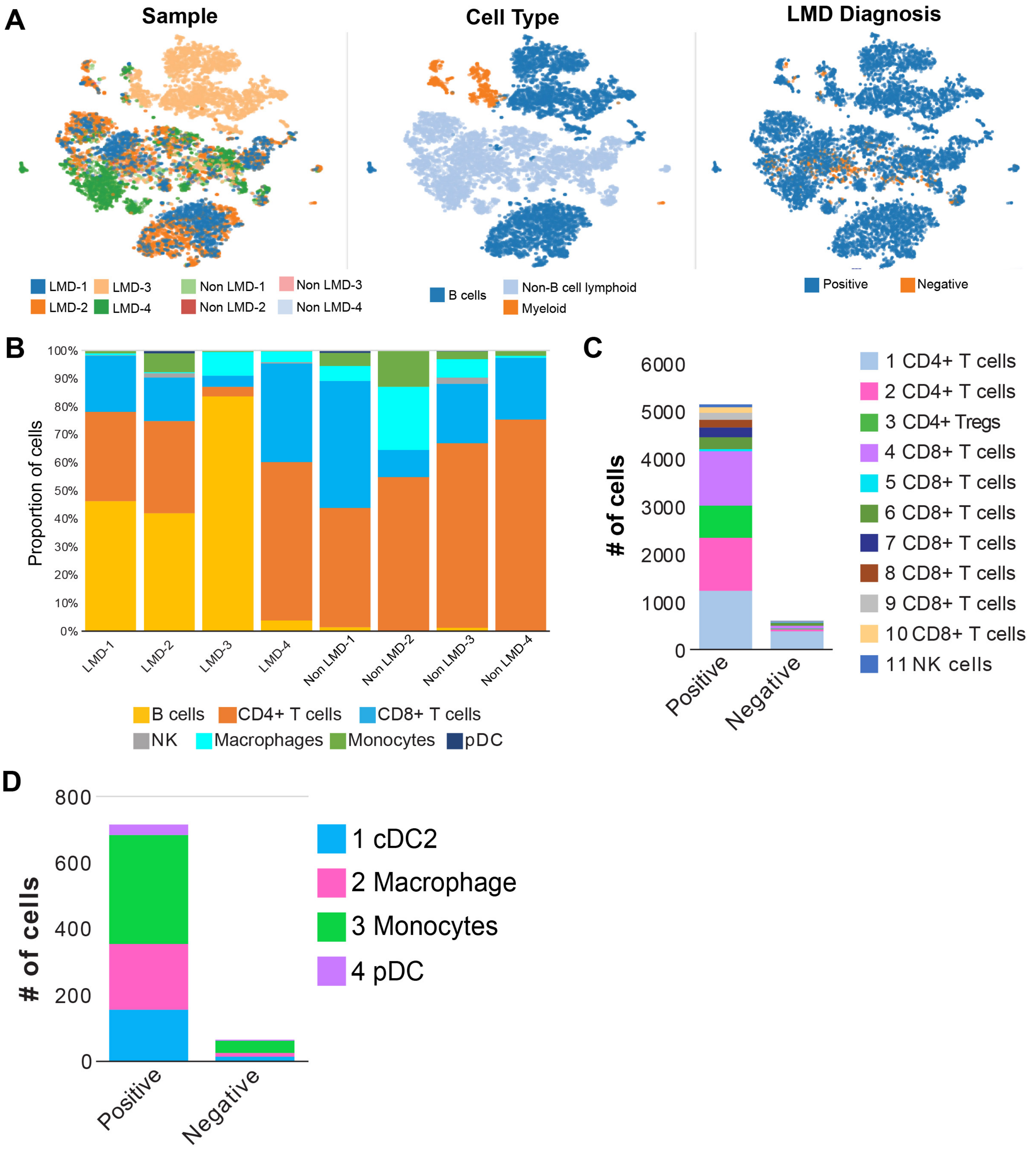


**Supplemental Figure 2. A.** t-SNE plots showing the clustering of the cells based on the sample of origin, cell type, and LMD diagnosis. **B.** Bar graphs showing the proportion of various cell types found in each sample. **C.** Bar graph showing the number of cells from various T cell subpopulations in each sample. **D.** Bar graph showing the number of cells from various myeloid cell subpopulations in each sample.

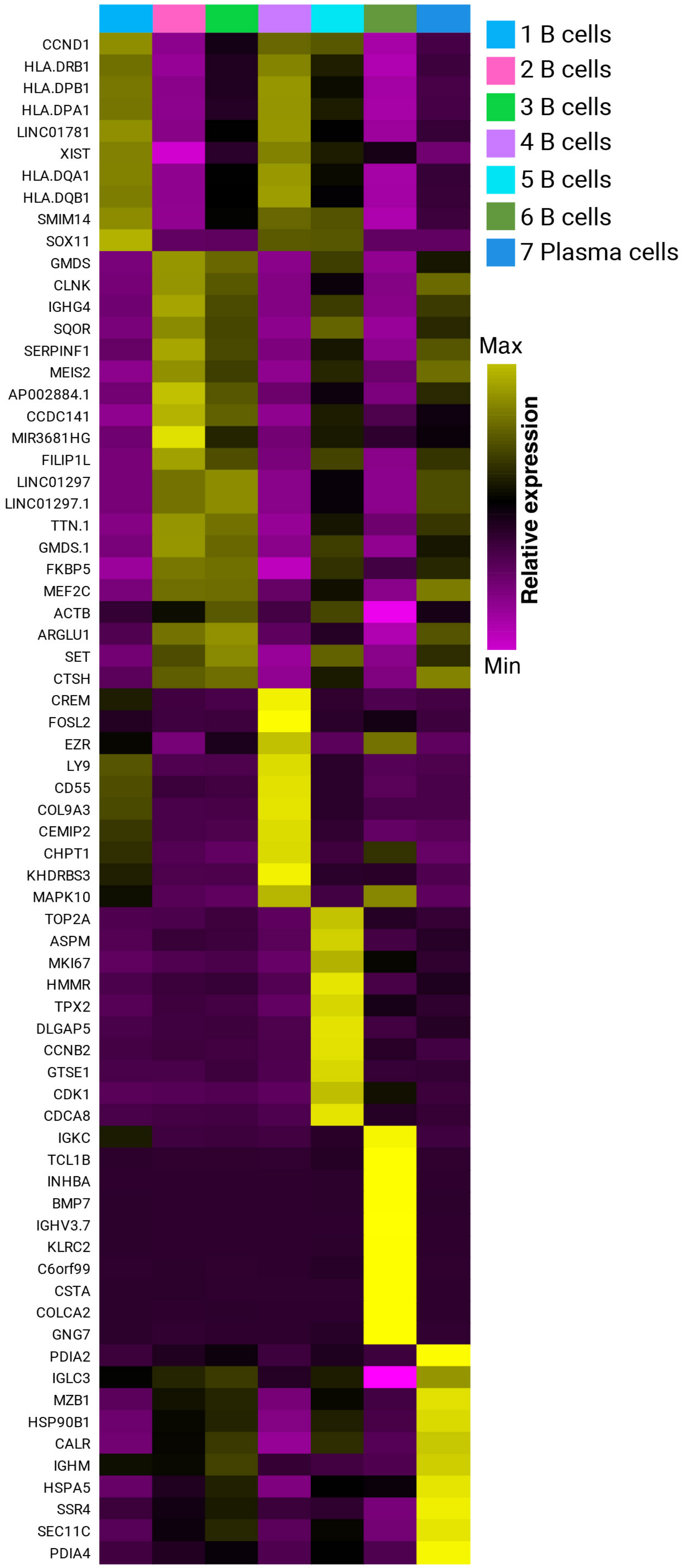


**Supplemental Figure 3.** Heatmap showing the relative expression of markers associated with each tumor (B cell) cluster.

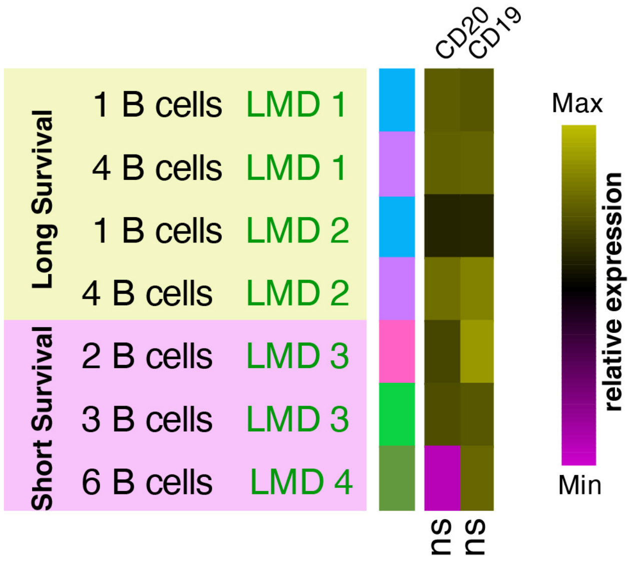


**Supplemental Figure 4.** Heatmap showing the relative expression of CD19 and CD20 in B cell clusters found in CSF of patients with long vs short survival. Differences in expression of CD19 and CD20 were both not significant (ns).

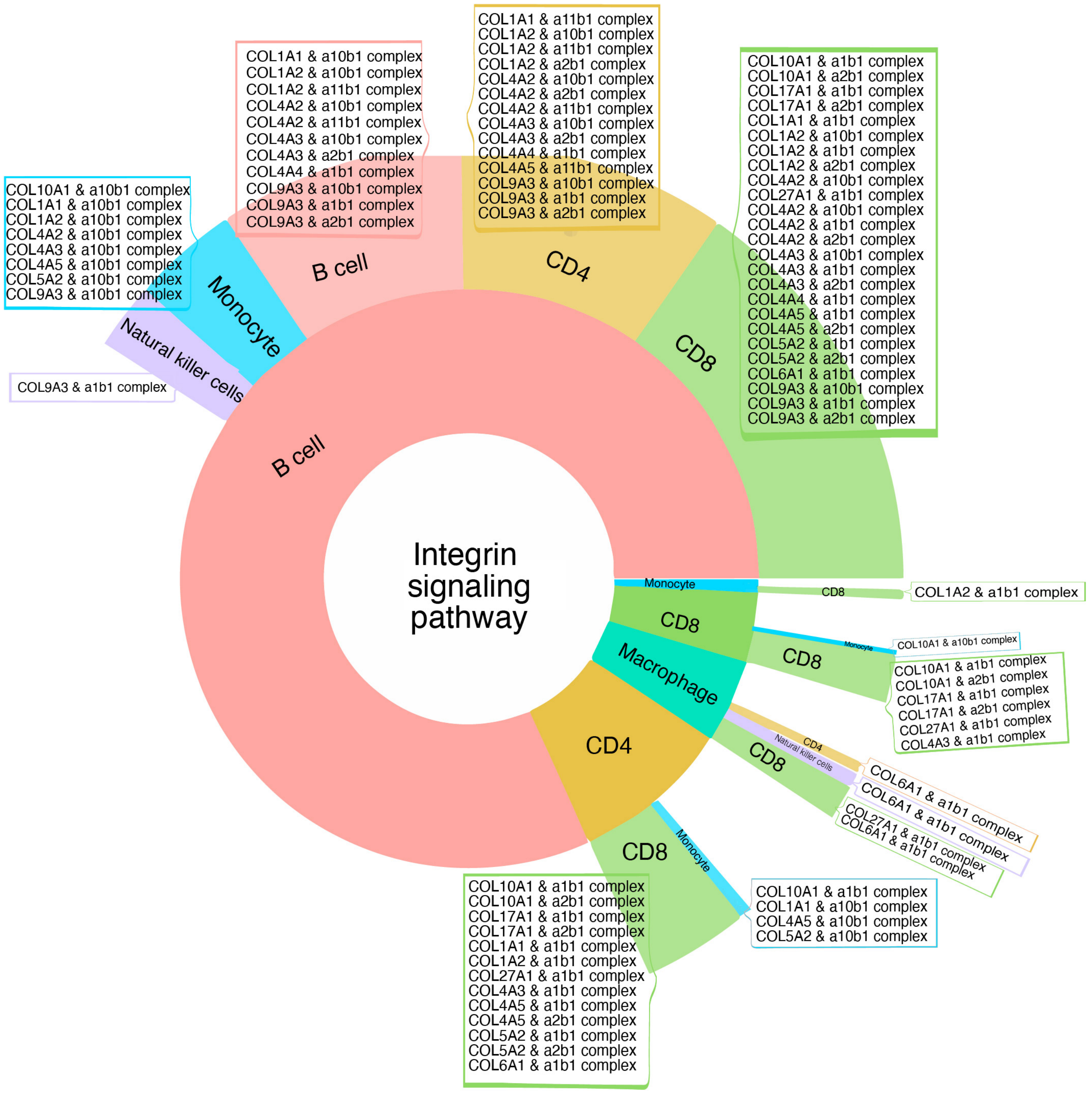


**Supplemental Figure 5.** Sunburst plot for Integrine signaling pathways as the most significant functional term based on only the unique ligand-receptor in patients with long survival. The width of each section represents the relative fraction of interactions (weighted by score) enriched in that cell type. Boxes show specific int-pairs enriched for the corresponding cluster pairs. Boxes were manually added to the figure from InterCellar downloaded files. All unique, condition-specific functional terms are shown in Supplemental Tables 5 & 6.

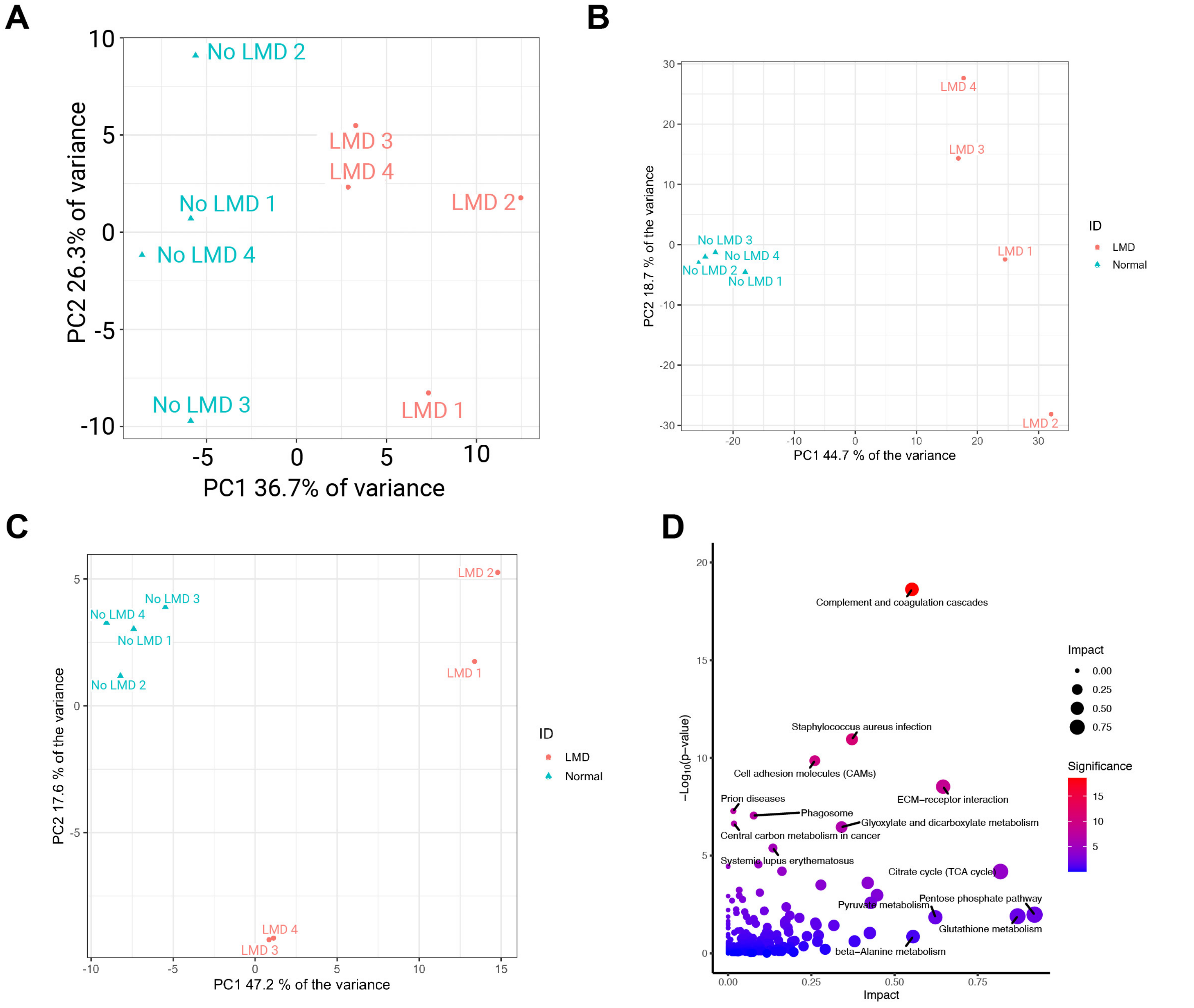


**Supplemental Figure 6. A.** Plot showing the PCA analysis of lipidomic profiling of CSF samples. **B.** Plot showing the PCA analysis of proteomic profiling of CSF samples. **C.** Plot showing the PCA analysis of metabolomic profiling of CSF samples. **D**. Plot showing results from integration of proteomics and metabolomic analysis of CSF from patients with LMD comparared to no LMD controls.

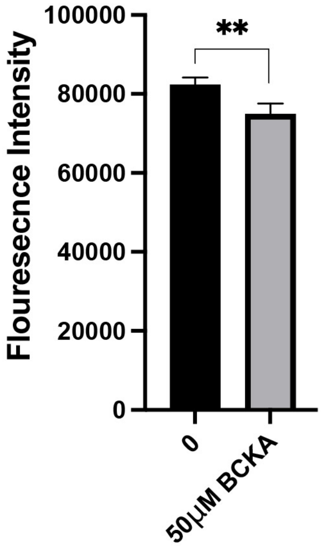


**Supplemental Figure 7**. Calcein AM Flouresence assay reflecting the vaibility of primary murine neurons under 50μ MBCKA in regular media for 7 days.

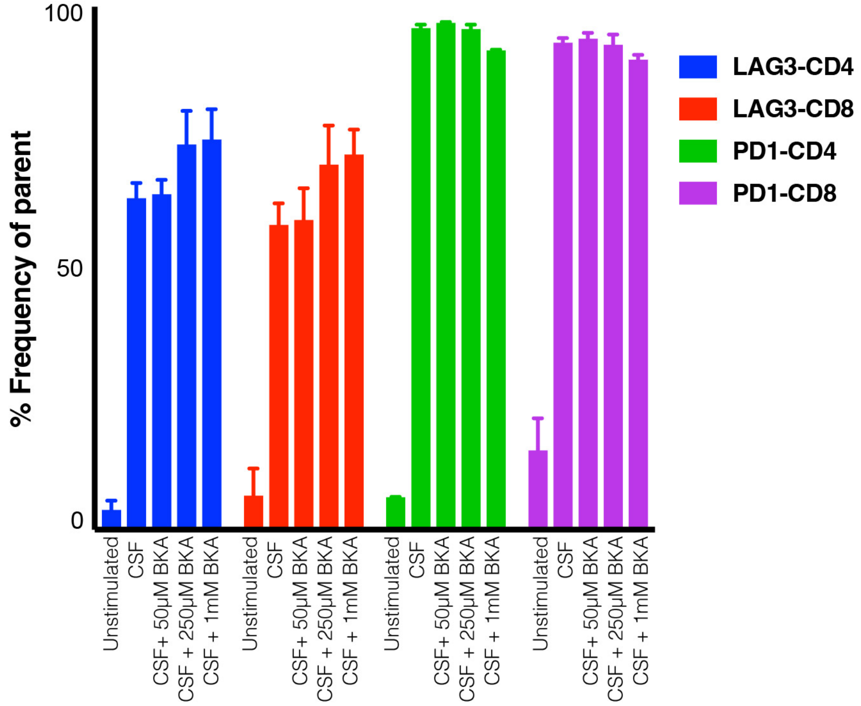


**Supplemental Figure 8.** Bar graph showing the expression of PD-1 and LAG3 on human CD4+ and CD8+ T cells in response to diiferent concentrations of BCKA in physiological CSF for 5 days. The frequency of PD1 and LAG3 positive cells were measured using flow cytometry. Data represent mean ± SD, data repeated using different human PBMCs.

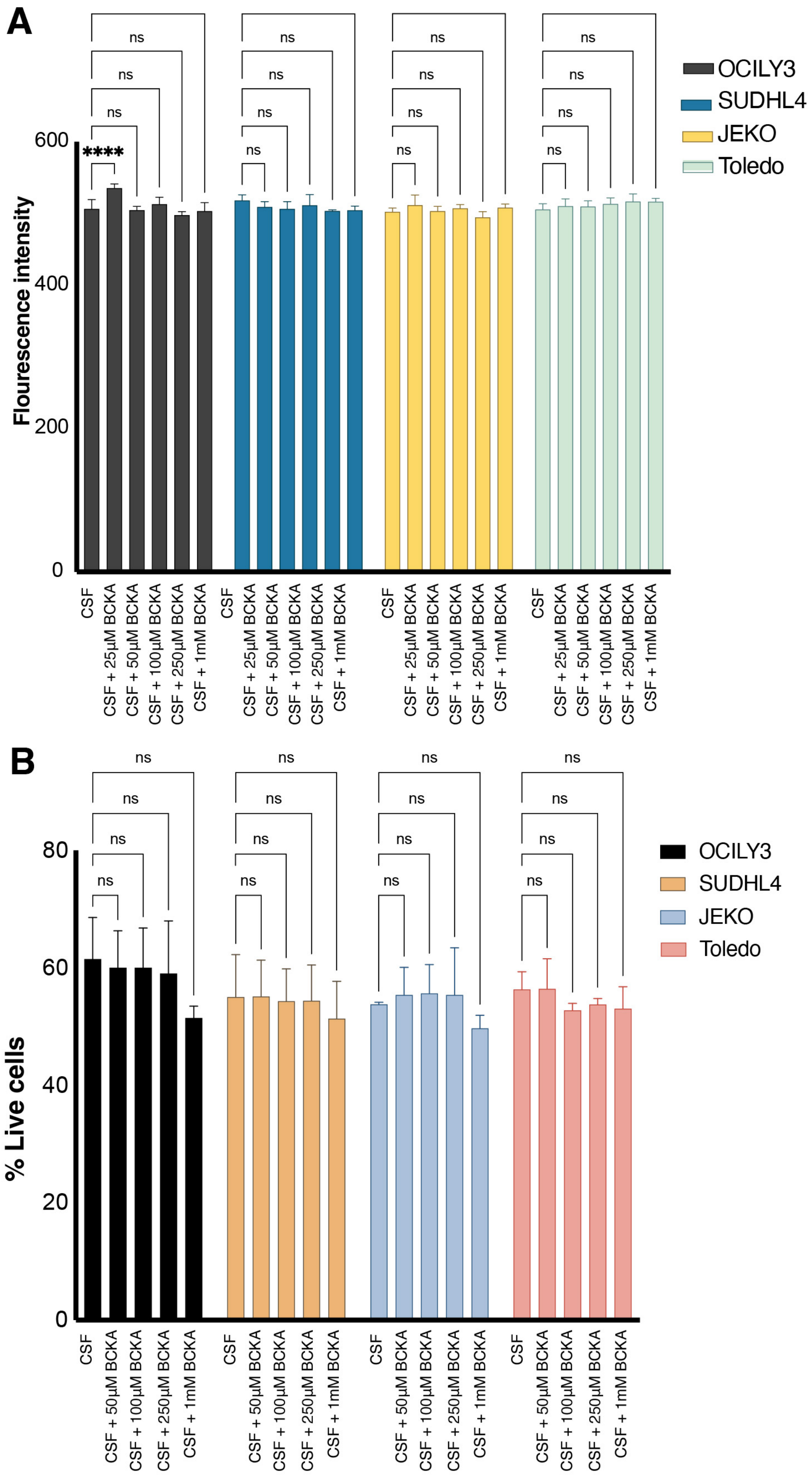


**Supplemental Figure 9**. **A**. CalceinAm assay on human lymphoma cell lines treated with increasing doses of BCKA at physiological ratio in physiological CSF for 48 hours. **B**. Proportion of live human lymphoma cell lines treated with increasing doses of BCKA at physiological ratio for 48 hours in physiological CSF based on excluding the propidium iodide postive cells (dead) using flow cytometery. Data represent mean ± SD, Statistical analysis was done using two-way ANOVA.****P <0.0001, ns=not significant

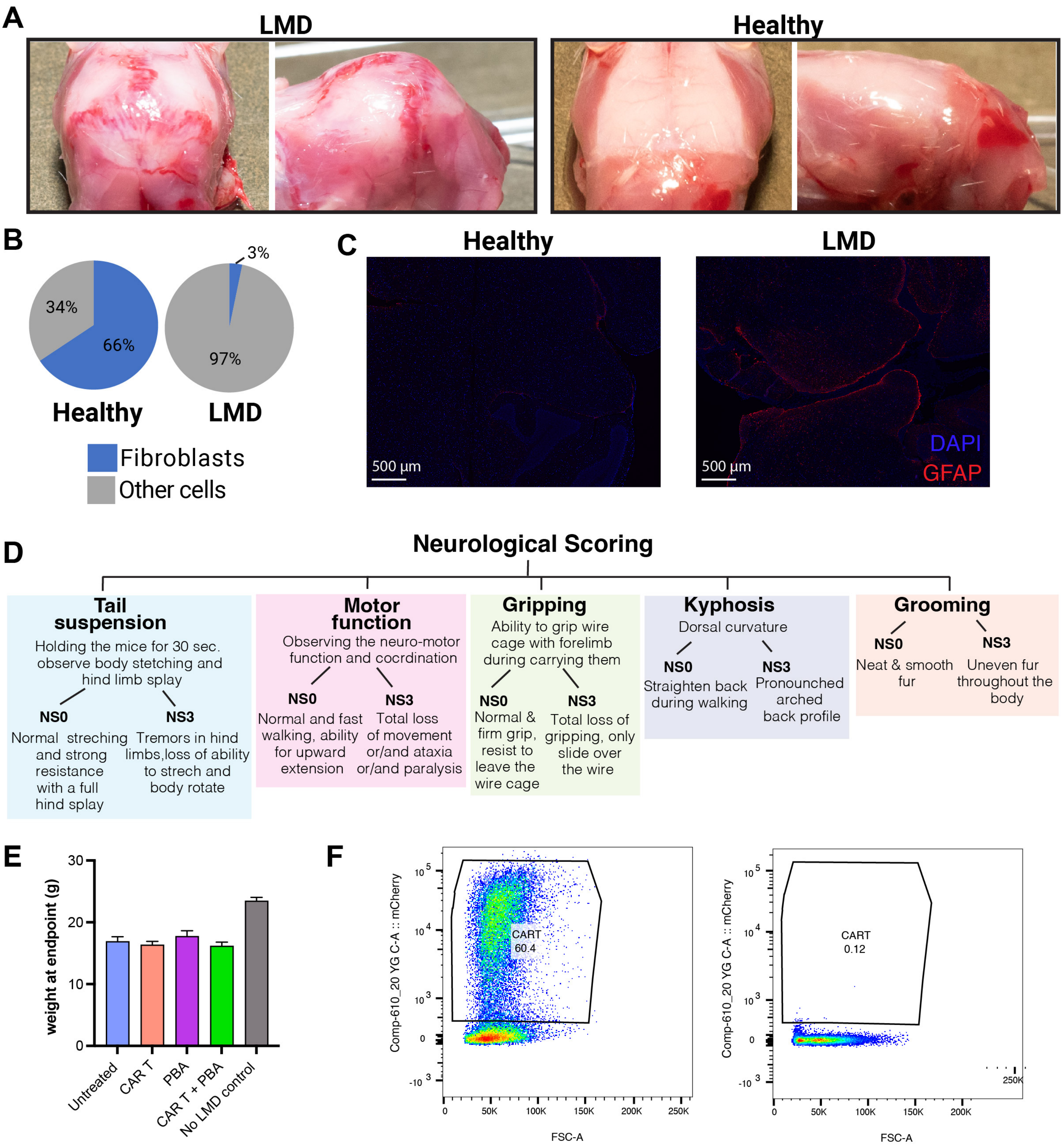


**Supplemental Figure 10. A.** Images showing the deformity in skull shape of the animals with lymphoma LMD. **B.** Proportion of fibroblast-like cells in healthy pia (PBS injection) versus LMD pia (A20 tumor injection) based on cell typing analysis from single cell RNAseq of the A20 animal models of LMD. **C.** Immunofluorescent staining for GFAP (Opal 690, red) and a DAPI nuclear stain (blue) on brain tissues of animals injected with A20 or PBS intrathecally. **D.** A schematic illustartion for the neuroscoring system in the LMD mouse mosel. **E.** Bar graph showing weights of animals at endpoint. **F.** Flow cytometry analysis for mouse antiCD19 CART-mCherry tagged postive cells compared to negative T cells at the day of mice injection.
